# Balancing of immune activation and suppression during phage infection

**DOI:** 10.64898/2026.06.04.730250

**Authors:** Iana Fedorova, Yourun Yue, Zirui Gao, Michelle Grunberg, Hao Wang, Junjie Li, Zhiyu Zang, Diego De Nault, Xilin Yang, Joseph P. Gerdt, Yue Feng, Joseph Bondy-Denomy

**Affiliations:** Department of Microbiology & Immunology, University of California, San Francisco, San Francisco, California, USA; State Key Laboratory of Green Biomanufacturing, College of Life Science and Technology, Beijing University of Chemical Technology, Beijing 100029, China; Department of Chemistry, Indiana University, Bloomington, IN 47405, USA

**Author notes:** co-first authors (Iana Fedorova, Yourun Yue). Corresponding authors; Joseph Bondy-Denomy, Yue Feng.

## Abstract

Signaling-based anti-bacteriophage systems such as CBASS and Thoeris synthesize infection-triggered nucleotide signals that activate anti-phage effectors^1,2^. However, the phage features sensed by these systems and the mechanisms phages use to evade signaling immunity remain poorly understood. Here, studying clinically relevant *Pseudomonas aeruginosa* phages from the *Migulavirinae* family^3^, we show that closely related phages encode subtle allelic variation in side tail fiber proteins that determine sensitivity to type II Thoeris. In parallel, these same phages encode an “anti-defense hotspot” that contains three adjacent genes that are each sufficient to facilitate phage evasion of both CBASS and Thoeris defenses, counter-balancing the activating proteins. Comparative analysis of this anti-signaling hotspot across the broader family of related N4-like phages uncovered a new Thoeris anti-defense (Tad) protein that sponges NAD-derived molecules (e.g. gcADPR) and exhibits sequence and structural similarity to a poorly characterized nucleotide-binding region of the human ryanodine receptor. Together, these findings reveal how the balance between immune activation and antagonism shifts phage outcomes and reveals a surprising similarity between a phage molecular sponge and an important human protein.

## Introduction

Bacteria encode diverse antiviral defense systems that protect against bacteriophage infection through mechanisms ranging from nucleic acid degradation to abortive infection and innate immune signaling^4^. Recent discoveries have revealed that many of the most widespread anti-phage systems rely on small molecule second messengers that activate defense effectors^5–7^. Among these, cyclic oligonucleotide-based anti-phage signaling systems (CBASS) and Thoeris have emerged as paradigmatic examples of bacterial innate immunity. CBASS systems detect phage infection and synthesize cyclic nucleotide signals such as 3′,3′-cGAMP or cyclic tri-adenylates, which activate diverse effector proteins that prevent viral propagation^8–11^. Similarly, Thoeris systems use Toll/interleukin-1 receptor (TIR) domain proteins to generate effector-activating NAD-derived signaling molecules^12–15^. The mechanisms by which the synthases detect phage infection remain poorly understood, however. In type I Thoeris, the phage capsid has been implicated as a trigger^16^ but for other Thoeris clades, including type II, the molecular features sensed by TIR-domain proteins in Thoeris systems remain unknown.

Bacteriophages have evolved a growing arsenal of anti-defense proteins that antagonize signaling immunity. Many recently identified anti-defense proteins function as “signal sponges,” directly binding immune signaling molecules to prevent effector activation^17^. Examples include anti-CBASS protein 2 (Acb2), which was initially identified to sequester CBASS cyclic nucleotides^18,19^. Thoeris anti-defense (Tad) proteins^13^ and Sequestins^20^ bind gcADPR signals produced by Thoeris. It was later learned that Tad and Acb proteins can have both multiple nucleotide binding sites and broad signal binding abilities^12,21–23^. To counter sponge-like proteins, hosts encode Panoptes systems that produce decoy cyclic nucleotides that suppress an effector. Sequestration or cleavage of these nucleotides by phage proteins de-represses the effector and halts phage propagation^24,25^. These findings suggest that signaling immunity drives intense evolutionary conflict between bacterial hosts and phages, yet the diversity, organization, and mechanisms of phage-encoded anti-signaling proteins remain incompletely explored.

Here, we make use of the N4-like bacteriophages within the *Migulavirinae* subfamily^3,26^, which infect *Pseudomonas aeruginosa,* to study anti-phage nucleotide signaling. Members of this phage family have been used clinically in humans^27^, and our previous work demonstrated that Lit1, another *Migulavirinae* phage, has its replication restricted by type II Thoeris in a multi-drug resistant clinical strain^28^. Therefore, understanding how these phages activate and/or resist signaling-based immunity is a pressing question. In this work we reveal that phage infection outcomes are governed by the balance between immune activation strength and layered anti-signaling mechanisms. Closely related *Migulavirinae* phages encode variations in side tail fiber proteins and their chaperones that dictate sensitivity to type II Thoeris, implicating phage-host adsorption-associated structures as determinants of immune sensing. In parallel, we identify a conserved anti-signaling hotspot that encodes multiple known and novel nucleotide sponges and broad evasion of signaling (*bes*) genes that reduce phage sensitivity to both CBASS and Thoeris defenses. Upon examining this compact genomic locus across diverse N4-like phages, we observe a bona fide anti-defense locus that contains homologs of many known nucleotide sponges. In this locus, we uncover a novel gcADPR sponge protein with surprising sequence and structural similarity to a poorly characterized nucleotide-binding region of the mammalian ryanodine receptor. Together, our findings reveal how phages simultaneously balance immune activation and immune inhibition and establish N4-like phages as a rich reservoir of anti-signaling mechanisms that illuminate fundamental principles of host-virus conflict.

## Results

### Viruses of the *Migulavirinae* subfamily display varying levels of sensitivity to Thoeris II immunity

Type II Thoeris is a two-gene phage defense system with a TIR-domain containing ThsB, proposed to sense phage infection through an unknown mechanism and generate Histidine-ADP ribose (His-ADPR) signals^12,14^. His-ADPR binds to a macrodomain-TM fusion protein, ThsA, which enacts a poorly characterized anti-phage mechanism (Fig. 1a). In a clinical *P. aeruginosa* isolate (MRSN11538^29^) the presence of type II Thoeris restricts the replication of the Lit1 bacteriophage (NC_013692, Fig. 1b)^30^ and dramatically reduces its burst size in a single round of infection (Fig. 1c).

**Figure 1.**
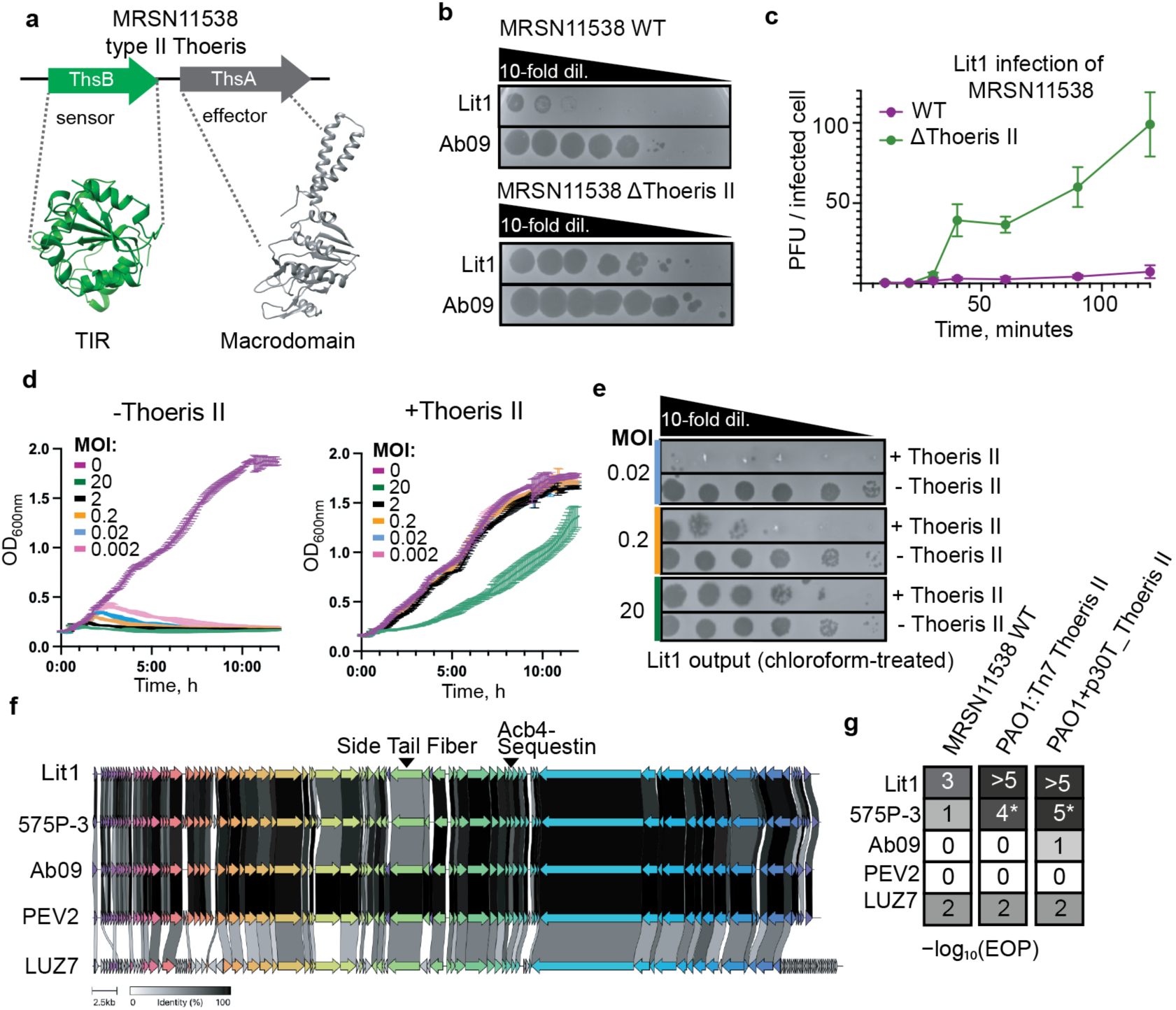
*Migulavirinae* phages are targeted by Thoeris Type II with different efficacy. **a**, A schematic of the MRSN11538 type II Thoeris locus with AlphaFold models of ThsA and ThsB proteins. **b**, Spot titration plaque assay of two indicated phages on bacterial lawns. Phage were titrated in 10-fold serial dilutions. **c**, Phage burst assay detecting released viral progeny over time in WT MRSN11538 (purple) or Δtype II Thoeris MRSN11538 (green). **d**, Growth curves measuring OD600 in liquid culture of PAO1 strains with (+Thoeris II) or without (-Thoeris II) type II Thoeris integrated in PAO1 chromosome during infection with different multiplicities of infection (MOI) of Lit1 phage. **e**, Output Lit1 phage production from the liquid cultures with (+Thoeris) or without (-Thoeris) defense presented in panel (d) after 12 hours of liquid culture infection. Output phage were titrated on PAO1:Tn7 empty lawns. **f**, Comparative bioinformatic alignment of *Migulavirinae* phages genomic sequences. Orthologous genes are indicated as arrows with the same colors. Alignment was generated with Clinker^32^. **g**, Efficiency of plating (EOP) of members of *Migulavirinae* phage family by type II Thoeris expressed endogenously from MRSN11538, integrated in the PAO1 genome (PAO1:Tn7 Thoeris II), or from a high copy plasmid (p30T). EOP was calculated as the phage titer on +Thoeris II cells divided by the phage titer on −Thoeris II cells.

Type II Thoeris (*thsB-thsA*) expressed from the chromosome of a phage-sensitive laboratory strain that lacks most phage defenses (PAO1) is sufficient to also prevent Lit1 replication (Fig. 1d): cells carrying type II Thoeris survive in liquid culture when MOI is low, while the defense is overwhelmed by Lit1 at high MOI, resulting in decreased optical density due to phage production (Fig. 1d,e). LC-MS measurement of His-ADPR signal in cells upon Lit1 infection shows that *P. aeruginosa* type II Thoeris produces His-ADPR signal in response to infection (Extended Data Fig. 1). To determine if the ThsA effector protein functions similarly to those described in recent reports^12,14^, we solved the crystal structure of the macro domain of MRSN11538 ThsA in the apo state and bound to His-ADPR (Extended Data Table 1). In the crystal it forms a dimer independently of His-ADPR presence (Extended Data Fig. 2a). The dimeric state of the apo protein was also confirmed by SEC-MALS (Size Exclusion Chromatography-Multi-Angle Light Scattering, Extended Data Fig. 2b). Comparison between the two structures shows that several of the residues surrounding the nucleotide signal display marked conformational changes (Extended Data Fig. 2c,d). Mutations in the ThsA macrodomain His-ADPR binding site abolish phage defense (Extended Data Fig. 2c and 3).

Interestingly, phage Ab09 (NC_024140), a Lit1-related (93% nucleotide identity) phage from within the *Migulavirinae* family, was not well-targeted by type II Thoeris in MRSN11538, with only a slight reduction in Ab09 plaque size observed (Fig. 1b). Due to the differences in Lit1 and Ab09 susceptibility to the system, we challenged type II Thoeris with additional bacteriophages from this family, phages Luz7, 575P-3 and PEV2. Comparative bioinformatic analyses showed that these phages are highly similar in their gene content, with LUZ7 (NC_013691) as the most divergent member (Fig. 1f). Lit1 and Luz7 are targeted by Thoeris II in MRSN11538, while Ab09, PEV2 (NC_031063), and 575P-3 (vB-Pae575P-3, NC_070865)^31^ are resistant (Fig. 1g). Induction of type II Thoeris expression, either from the chromosome (Tn7) or multi-copy plasmid (p30T) in PAO1, increased the strength of 575P-3 targeting (Fig. 1g, Extended Data Fig. 4). We further divide *Migulavirinae* phages into “strongly targeted” (Lit1), “moderately targeted” (575P-3), and “non-targeted” (Ab09, PEV2) by type II Thoeris.

### *Migulavirinae* side tail fibers and their chaperones determine type II Thoeris sensitivity

Phages that resist type II Thoeris (Ab09 and PEV2) may possess inhibitory factors, have variations in sensed phage components, or a combination thereof. To understand which Ab09 genes are required for immunity to type II Thoeris, we performed Lit1 and Ab09 phage hybridization. MRSN11538 cells naturally encoding the wild-type type II Thoeris system that targets Lit1 were transformed with a plasmid expressing the type I-C CRISPR-Cas3 system^33^ with crRNA uniquely targeting Ab09 (Ab09 *gp80*). This strain is resistant to parental Lit1 and Ab09, and therefore selected for hybrid phages that escape both systems. Co-infections gave rise to hybrid Lit1-Ab09 plaques, which were isolated and sequenced (Fig. 2a). The hybrids exhibited complete resistance to type II Thoeris in MRSN11538 and complete or partial resistance to Thoeris immunity in PAO1:Tn7 Thoeris II strain, where defense is stronger (Extended Data Fig. 5). Sequencing of the Lit1/Ab09 hybrids revealed highly variable patterns of Ab09- and Lit1-derived regions, but all hybrids shared three commonly acquired Ab09 sequences, designated regions 1, 2, and 3 in Fig. 2b.

**Figure 2.**
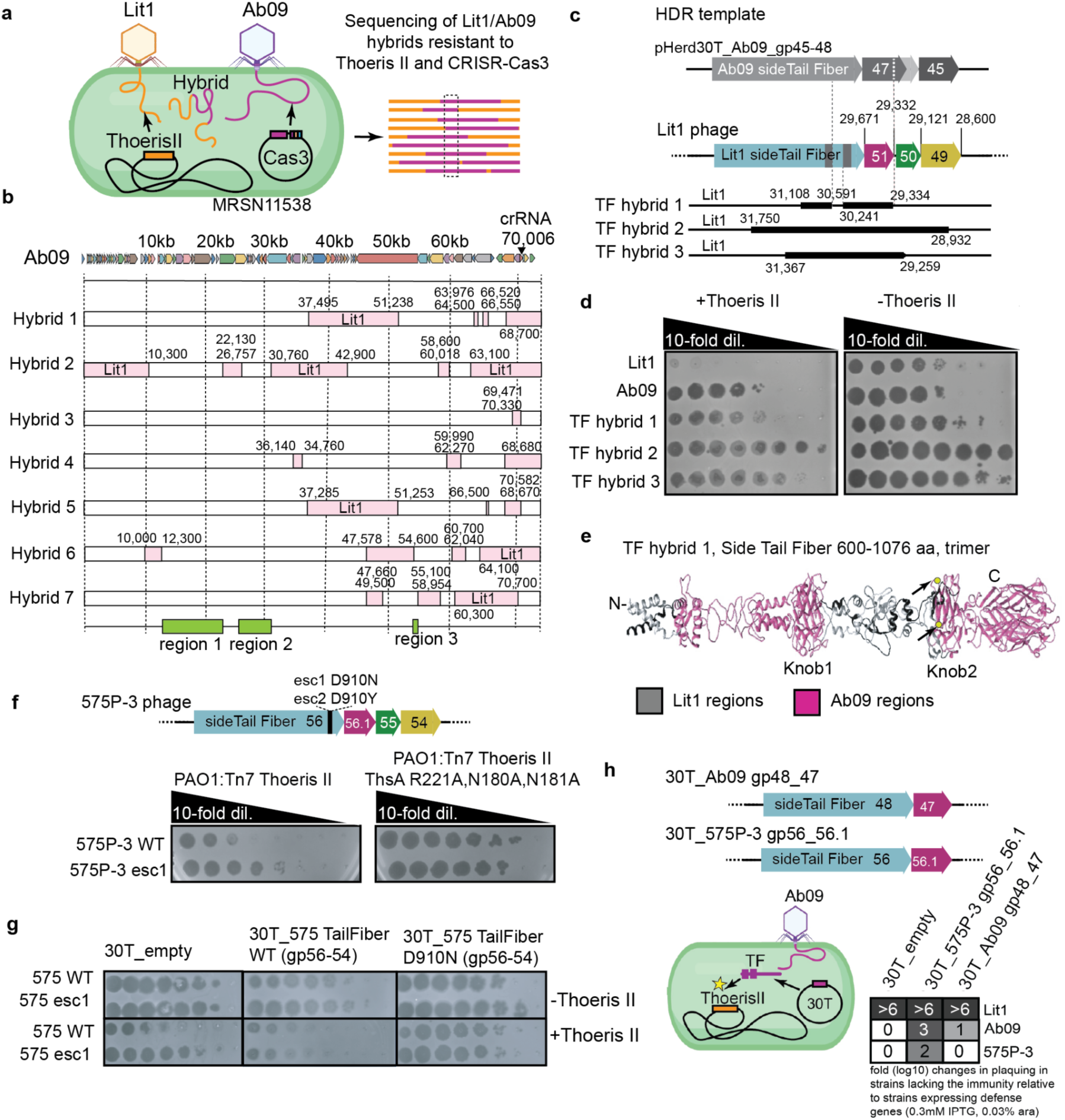
Side tail fiber and chaperone alleles determine type II Thoeris sensitivity. **a**, A scheme of the phage hybridization assay. Lit1 (type II Thoeris targeted) and Ab09 (CRISPR-Cas3 targeted) phages infected the MRSN11538 strain carrying both defenses. **b**, Sequences of type II Thoeris-resistant Lit1/Ab09 hybrids. Regions derived from Ab09 and Lit1 are shown as white and pink boxes, respectively. **c**, Schematic showing type II Thoeris-resistant Lit1 hybrids, produced through recombination with an HDR template carrying the indicated Ab09 genes. The coordinates of genes in Lit1 phage are shown. Positions of knob domains are shown as grey rectangles. **d**, Spot titration plaque assay of Lit1 side tail fiber hybrids on bacterial lawns. Phage was titrated in 10-fold serial dilutions. **e**, AlphaFold model of the C-terminal end of the Lit1 side tail fiber, 600-1076 amino acids, trimer, showing regions derived from Ab09 (pink) in type II Thoeris-resistant Hybrid1. Arrows show positions that correspond to D910 in 575P-3 phage side tail fiber (position of escaper mutation). **f**, 575P-3 type II Thoeris escapers. Above - scheme showing a position of escape mutation, below - spot titration plaque assay of 575P-3 wild type (WT) and 575P-3 gp56 D910N (esc1) on bacterial lawns. **g**, Spot titration plaque assay of 575P-3 wild type WT and 575P-3 Esc1 on bacterial lawns overexpressing type II Thoeris and 575P-3 side tail fiber (WT and D910N) with adjacent genes. Phage was titrated in 10-fold serial dilutions. **h**, Overexpression of 575P-3 side tail fiber with its putative chaperone. Above - a schematic of vectors used for the overexpression of side tail fibers and adjacent genes. Below - fold (log10) changes in plaquing in strains lacking immunity relative to strains with type II Thoeris.

In the Lit1-Ab09 hybrid phages, Region 2 shows the greatest sequence divergence between Lit1 and Ab09, and encodes structural proteins, specifically side tail fibers, which we considered to be potential ThsB activators. To query whether the acquisition of the Ab09 side tail fiber and associated genes from region 2 bestow type II Thoeris resistance upon Lit1, we provided Ab09 *gp48* (side tail fiber), *gp47*, *gp46*, and *gp45* as a single homology-directed repair (HDR) template for recombination into the Lit1 genome (Fig. 2c). The resulting phage lysate, a mix of wild-type Lit1 and various hybrids that acquired regions of the HDR template, was selected on type II Thoeris-expressing cells, on which Lit1 does not form spontaneous escapers. Rare surviving plaques revealed phages that had acquired various regions of the Ab09 side tail fiber locus (i.e. derived from region 2), confirming the causality of this locus in enabling Thoeris escape. Hybrid 1 acquired the shortest region from Ab09 compared to the other sequenced hybrids, which included the 3′ end of the side tail fiber gene (gp48) and its adjacent gene (gp47, Fig. 2c,d). Prediction of Lit1 side tail fiber structure using AlphaFold showed that the replacement of the C-terminal region of the protein with the Ab09 variant in the Thoeris-resistant Hybrid 1 occurred primarily in-and-around the knob domain region of the side tail fiber (Fig. 2e), suggesting this region from Ab09 can reverse type II Thoeris sensitivity.

Supporting the importance of the side tail fiber knob domain in type II Thoeris sensitivity, we also isolated two spontaneous phage 575P-3 escapers on type II Thoeris-expressing PAO1, which carried a D910N or D910Y mutation in the Knob2 region of the 575P-3 side tail fiber (Fig. 2f). Over-expression of the 575P-3 side tail fiber gene alone did not efficiently complement these mutations (i.e. escape phages were not targeted), suggesting that this gene is insufficient to enhance type II Thoeris activity (Extended Data Fig. 6a). As tail fiber protein folding often requires chaperones encoded downstream^34–37^, we thus added gp56.1 (Ab09 gp47 ortholog, unannotated in 575P-3 NCBI genome), gp55, and gp54 to the construct. Expression of WT 575P_gp56-54 increased the type II Thoeris targeting of WT 575P-3 and 575P-3 escaper 1, while the same construct with the D910N mutation decreased the targeting and helped WT 575P-3 to escape (Fig. 2g). Further minimization of this construct demonstrated that expression of just the 575P-3 side tail fiber and its adjacent putative chaperone (gp56 and gp56.1* for 575P-3) led to an increase in type II Thoeris targeting of both Ab09 and 575P-3. This activation phenotype (i.e. increased phage targeting) is dependent on the presence of the His-ADPR binding site in ThsA, demonstrating that canonical type II Thoeris activity is indeed being stimulated (Extended Data Fig. 6b). With the homologous genes from Ab09, a naturally non-targeted phage, apparent type II Thoeris activation was less potent (Fig. 2h, Extended Data Fig. 7). PEV2, which is identical to Ab09 through this region, is also not targeted by type II Thoeris. The 575P-3 side tail fiber amino acid sequence is diverged from those of Ab09 and Lit1, but the proteins exhibit high structural similarity (Extended Data Fig. 8). Modeled co-folding of the 575P-3 side tail fiber gp56 proteins with gp56.1 and gp55 supports their potential role as tail fiber chaperones, which are similar to F2 pyocins chaperones PyoF14 and PyoF15^38^ (Extended Data Fig. 9). We propose that the knob domains of side tail fibers of *Migulavirinae* phages, properly folded by, or in complex with, its chaperone, determines the sensitivity to type II Thoeris.

### Discovery of a hot spot of phage anti-signaling genes

We next queried whether the tail fiber locus, on its own, is sufficient to explain Ab09 resistance to type II Thoeris or whether the phage additionally possesses inhibitory factors against this or other signaling systems. In testing type II-A CBASS (3′,3′-cGAMP), type III-C CBASS (cAAA) or type I Thoeris (gcADPR), we observed that Ab09 is completely resistant to all systems (Fig. 3a). To explain this broad signaling defense evasion, we examined Ab09 for the presence of known anti-defense genes and found *gp59*, encoding a fusion protein of Acb4 (Type II-A CBASS antagonist^39^) and Sequestin (type I Thoeris antagonist^20^). Over-expression of Ab09 *gp59* showed strong anti-Thoeris type I activity and partial type II-A CBASS antagonism (Extended Data Fig. 10). No inhibition of type II Thoeris was observed. Acb4-Sequestin very tightly binds 1′′,3′-gcADPR *in vitro* and more weakly to 3′,3′-cGAMP (Fig. 3b), explaining how it inhibits both systems. SEC-MALS showed that the full-length protein, Acb4 and Sequestin domain forms a tetramer, tetramer and dimer, respectively (Extended Data Fig. 11). We also solved the crystal structure of the Sequestin domain (Extended Data Table 1), which possesses a similar fold to what was previously predicted^20^, i.e., an antiparallel dimer with two symmetrical pockets in the interface between protomers (Fig. 3c). Despite the strength of antagonism of the type I Thoeris system, deletion of *gp59* from Ab09 surprisingly yielded a phage that remained resistant to all signaling defenses (Fig. 3d), suggesting that Ab09 may possess additional anti-signaling genes, possibly encoded nearby.

**Figure 3.**
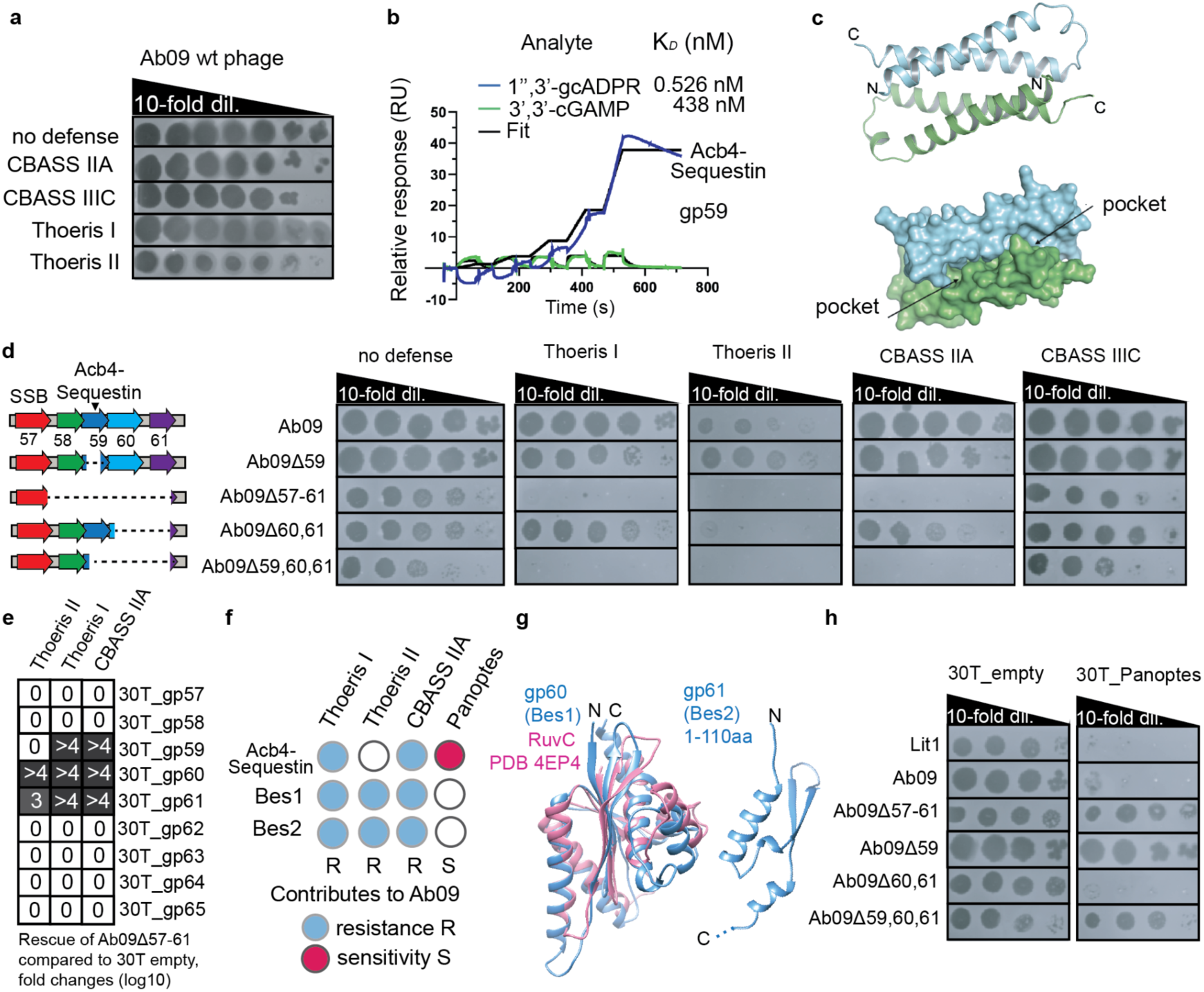
The combined activity of three non-essential genes helps Ab09 to protect against various signaling-based defenses. **a**, A spot titration plaque assay of Ab09 wild type (wt) phage on bacterial lawns. Phage was titrated in 10-fold serial dilutions. **b**, Overlay of sensorgrams from SPR experiments used to determine kinetics of gp59 binding to 1′′,3′-gcADPR and 3′,3′cGAMP. Data were fitted with a model describing one-site binding for the ligands (black lines) and (black lines) **c**, Overall structure of the Sequestin domain of Acb4-Sequestin shown in cartoon model and surface model, respectively. **d**, Spot titration plaque assay of phage Ab09 and various deletion mutants on indicated bacterial lawns. Left, a scheme of the deletions in Ab09. **e**, Efficiency of rescue of Ab09Δ57–61 phage on bacterial lawns carrying corresponding defenses in the presence of indicated Ab09 genes, encoded on plasmids (30T), compared to empty vector. **f**, A scheme showing a contribution of Ab09 anti-signaling proteins to resistance or sensitivity to indicated defense systems. **g**, AlphaFold model of gp60 (Bes1) (blue) and RuvC protein (PDB 4EP4) (pink) aligned to each other (RMSD 2.9 Å) and AlphaFold model of gp61 (Bes2). **h**, Spot titration plaque assay of indicated phages on bacterial lawns overexpressing the Panoptes defense system. Phage was titrated in 10-fold serial dilutions.

To determine whether the *gp59* locus is an anti-defense island^40–42^, we deleted *gp59* and surrounding genes (Fig. 3d). The deletion, generated without prescribed boundaries by Cas3 (see methods), yielded phage Ab09*Δgp57-61* lacking 51 bp in the 3’ end of *gp57* (a putative SSB), all of *gp58* (unknown function), *gp59* (Acb4-Sequestin), *gp60* (putative RuvC domain), and most of *gp61* (unknown function) (41,096-42,899 positions) (Fig. 3d). This mutant phage surprisingly became sensitive to type II Thoeris, type II-A CBASS, and type I Thoeris (Fig. 3d). Targeting of Ab09*Δgp57-61* is dependent on the presence of active immune effectors (Extended Data Fig. 12). This mutant phage had slightly reduced plaque size but generally replicated well in the absence of immunity and remained resistant to type III-C CBASS. Complementation with single ORFs surprisingly revealed that *gp59*, *gp60*, and *gp61* all individually reversed defense activity of multiple systems (Fig.3e, Extended Data Fig. 13), suggesting functional redundancy. Indeed, a precise deletion of all three genes, *gp59*, *gp60*, and *gp61*, rendered Ab09 susceptible to these defenses, whereas single (Ab09*Δ59)* or double (Ab09*Δ60,61)* deletions remained type I Thoeris-resistant and partially resistant to type II-A CBASS (Fig. 3d). Consistent with the observation of *gp60* and *gp61* anti-Thoeris type II activity, phage Ab09*Δ60,61* failed to propagate on type II Thoeris (Fig. 3d). Based on these results, we conclude that Ab09 bacteriophage uses the combined activities of *gp59 (acb4-sequestin)*, *gp60 (*here named *broad evasion of signaling 1, bes1)*, and *gp61 (bes2)* to evade type I Thoeris, type II-A CBASS, and type II Thoeris (Fig. 3f). AlphaFold predictions show that Bes1 is related to RuvC domains, while Bes2 lacks recognizable structural similarity to characterized proteins and is predicted with low confidence over most of its sequence (Fig. 3g). Moreover, this locus is generally conserved in the *Migulavirinae* phages, suggesting that the weakly Thoeris-activating tail fiber locus (e.g. Ab09, PEV2) does not “rise above” this type II Thoeris antagonism, while strongly activating phages (e.g. Lit1), do.

The Panoptes phage defense system uses decoy 2′,3′-cAA (2′,3′-c-di-AMP) signals, whose sequestration by phage-encoded sponges Acb2 or Tad1/2 activates immunity^24,25^. We asked whether activation of Panoptes could be a convenient bioassay to determine whether Bes1 and Bes2 bind/degrade cyclic di-nucleotides. Using the characterized *KP67* Panoptes system, we observed restriction of wild-type Lit1 and Ab09 phages, while the Ab09*Δgp57–61* mutant completely escapes Panoptes (Fig. 3h). Interestingly, deletion of just Ab09 *acb4-sequestin* leads to full recovery of the phage when infecting cells expressing Panoptes (Fig. 3h). This Acb4-Sequestin fusion binds 2′,3′-cAA signals *in vitro* and releases the signal upon inclusion of SDS, supporting its action as a sponge (Extended Data Fig. 14). This is consistent with recent work showing that *acb4* encoded by the N4 *E. coli* phage activates Panoptes^25^. Deletion of *bes1-2* did not reduce Panoptes activity against Ab09, suggesting that the anti-signaling mechanisms of these proteins do not impact 2′,3′-cAA signaling (Fig. 3h). These data, together with *in vitro* experiments with Bes1 and Bes2 that showed no apparent binding or cleavage of 3′,3′-cGAMP or His-ADPR (Extended Data Fig. 15), suggest that Bes1 and Bes2 have a novel mode of protecting phages from signaling, which avoids the cost of triggering Panoptes (Fig. 3f). In sum, this work demonstrates that immune signaling sensitivity within the *Migulavirinae* phage is governed by the balance between the strength of immune activation and mechanistically diversified anti-signaling mechanisms.

### N4-like phages infecting other hosts possess signaling inhibitors

The phage gene island described here in the *Migulavirinae* phages motivated us to ask if this region is conserved in related phages infecting other bacteria. We set the most conserved genes in the region: the virion RNA polymerase (*gp66* in Ab09) and the DNA primase-polymerase (*gp55* in Ab09) as the boundaries and searched based on amino acid sequence and predicted structural similarity using SeqHub/Gaia^43^. We found that most N4-related phages carry this hot spot of gene acquisition, which encodes highly diverse genes within it, including known signaling-based inhibitors such as Tad2, Tad3, Acb4, and Sequestin (Fig. 4). Genes of unknown function are incorporated between DNA primase-polymerase, virion RNA polymerase, and interspaced with phage single strand binding protein (SSB) and RecA homologs. Orthologs of Bes1, all containing a predicted RuvC domain, are found in most of the analyzed genomic regions. Genes encoding proteins with structural similarity to Bes2 are also observed (Fig. 4).

**Figure 4.**
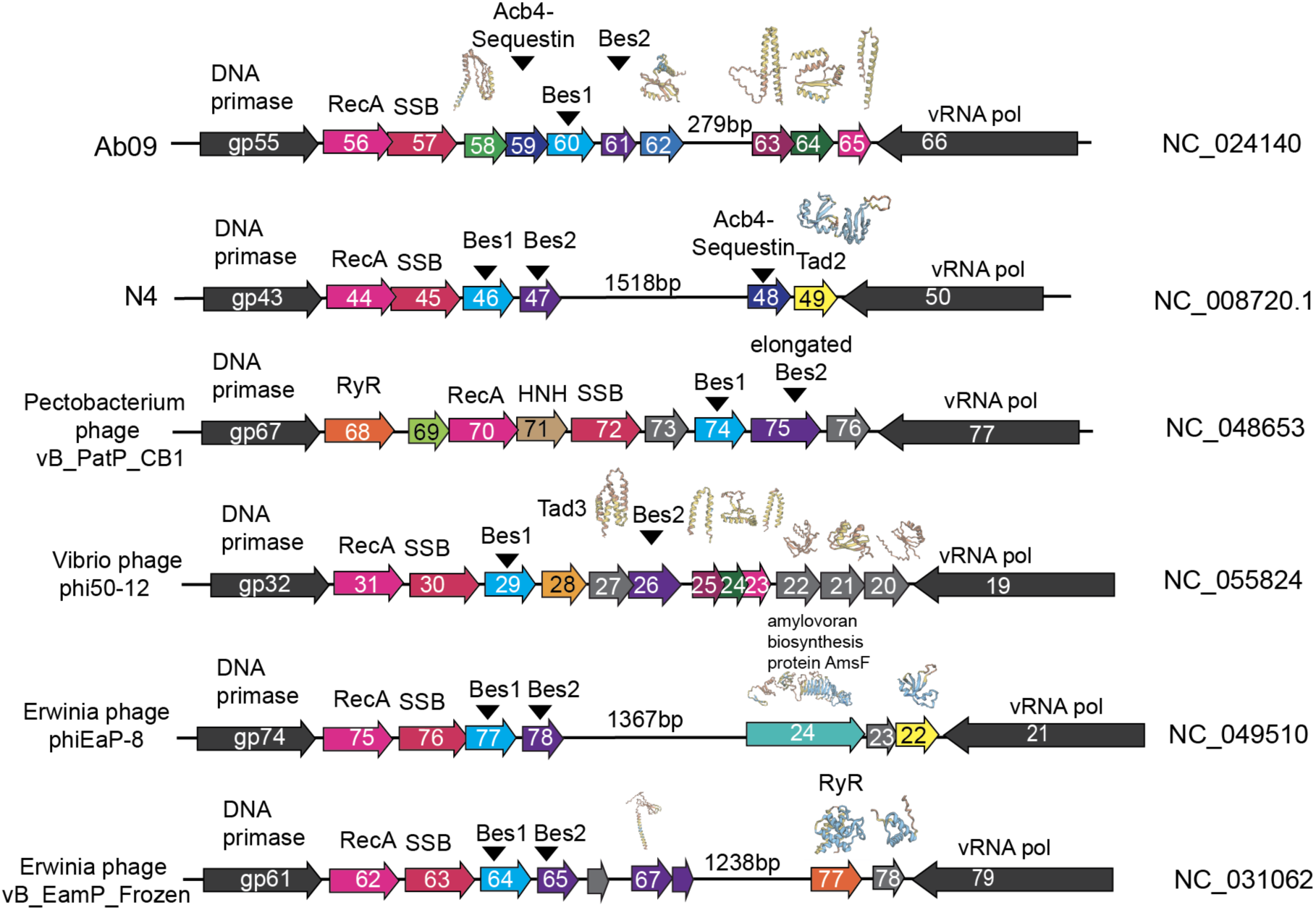
Organization of gene-acquisition hotspots in N4-related bacteriophages. Genes are shown as arrows, with matching colors indicating orthologous genes based on sequence or structural similarity. Structural predictions of selected ORFs are shown above the gene schematics.

### RyR-related Thoeris II inhibitor sequesters cADPR

Iterative expression of many genes in this putative anti-signaling hotspot revealed that *gp68* from *Pectobacterium* phage vB_PatP_CB1 (NC_048653) (Fig. 4) possessed robust anti-type I Thoeris activity and rescued both Ab09 *Δ57-61* and the unrelated type I Thoeris-sensitive model phage F10 (Fig. 5a). gp68 is encoded immediately adjacent to the conserved DNA primase and contains two distinct domains (Fig. 5b). The N-terminal domain constitutes a fold with unknown function, which is often encoded as a single ORF, while HHpred revealed that the C-terminal domain is highly similar to the Repeat12 domain (Extended Data Fig. 16) of the mammalian ryanodine receptors (RyR1-RyR3)^44,45^. This similarity extends to the primary sequence level, with 33% amino acid identity with a.a. 863-936 in human RYR3 (Fig. 5c). This receptor is responsible for essential biological processes in mammals, including the rapid release of Ca^2+^ from endoplasmic reticulum to the cytosol during muscle contraction. The function of the Repeat12 domain (shown in Fig. 5d) in the human ryanodine receptor is not fully understood, but it is both a drug and ATP binding site, proposed to play a role in metabolic sensing^46^. Splitting the phage protein into two fragments (gp68_N 1-86 a.a., gp68_C 65-183 a.a.) revealed that gp68_N had no apparent activity while gp68_C alone, the RyR-like region, is responsible for type I Thoeris inhibition (Fig. 5e). We now refer to gp68 as RyR Tad (Ryanodine receptor Thoeris anti-defense).

**Figure 5.**
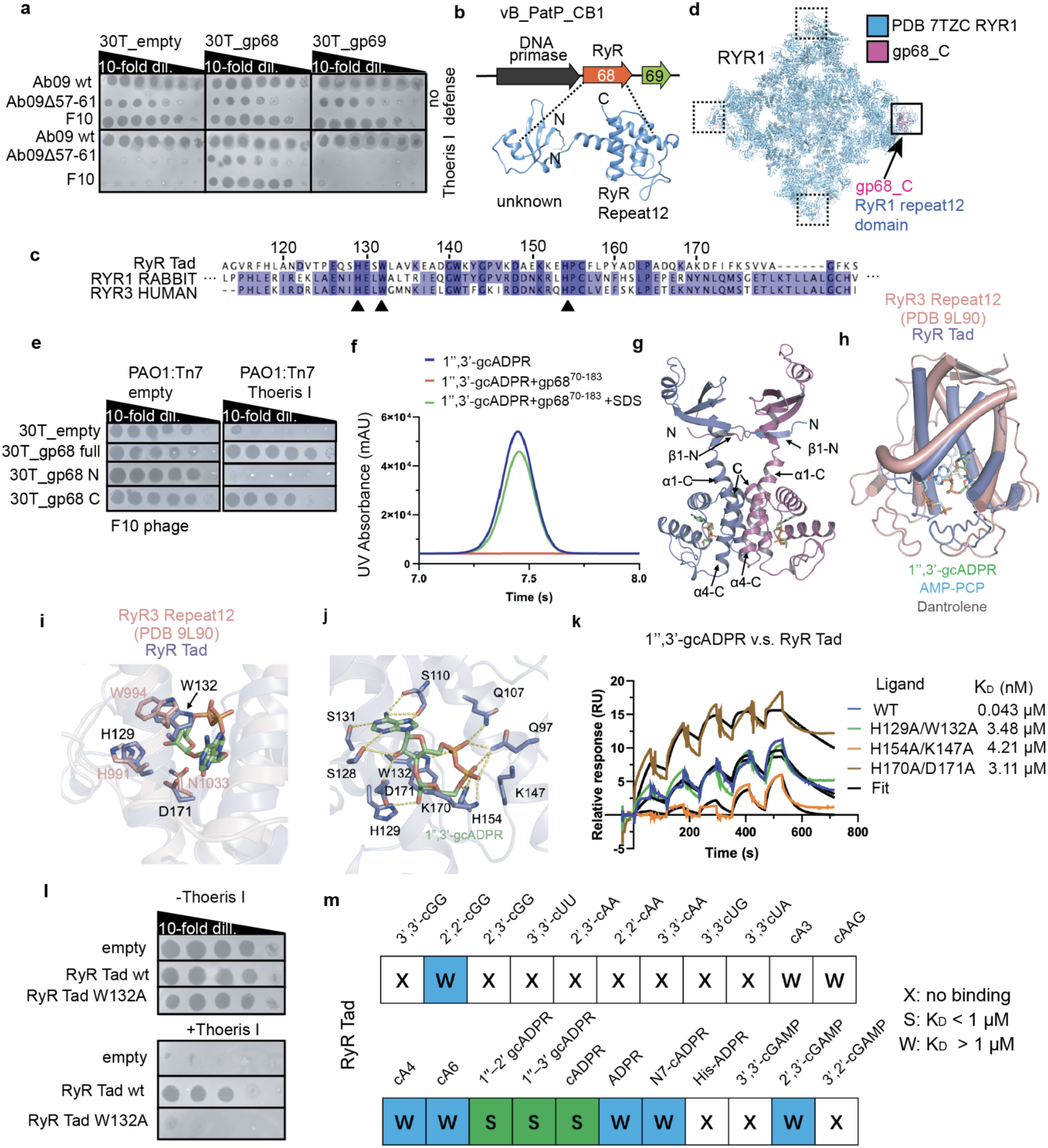
Phage RyR Tad is related to RyR repeats1,2 and possesses anti-Thoeris type I activity. **a**, Spot titration plaque assay of indicated phages on bacterial lawns with and without type I Thoeris defense, overexpressing vB_PatP_CB1 gp68 and gp69. Phage was titrated in 10-fold serial dilutions. **b**, Scheme of vB_PatP_CB1 phage gp68 carrying genome fragment. AlphaFold model of gp68 is shown. **c**, Sequence alignment of gp68 (RyR Tad) and Repeat12 domains of rabbit RYR1 and human RYR3 receptors (Uniprot IDs P11716 and Q15413-1). Arrowheads label residues that are required for binding to 1′′–3′ gcADPR binding. **d**, Structural alignment of AlphaFold model of gp68 C-terminal domain (pink) and RYR1 channel (blue) (PBD 7TZC). Positions of Repeat12 domains are shown in rectangles. **e**, Spot titration plaque assay of F10 phage on bacterial lawns with and without type I Thoeris defense, overexpressing N- and C-terminal domains of gp68 (1-86 and 65-183 aa) and its full-length version. Phage was titrated in 10-fold serial dilutions. **f**, The ability of gp68_C (70-183 aa) to bind and release 1′′,3′-gcADPR when treated with SDS was analyzed using HPLC. 1′′,3′-gcADPR standard was used as a control. The remaining nucleotides after incubation with gp68_C were tested. **g**, Overall structure of RyR Tad-1′′,3′-gcADPR complex. 1′′,3′-gcADPR is shown as sticks. β1-N, α1-C and α4-C represent β1 of the N-terminal domain, α1 and α4 of the C-terminal domain, respectively. **h**, Structural superimposition between RyR Tad C-terminal domain and the Repeat12 domain of human RYR3 (PDB: 9L90). The small molecules in the structure are shown as sticks. **i**, Structural superimposition between RyR Tad C-terminal domain and the Repeat12 domain of human RYR3 with three conserved residues involved in 1′′,3′-gcADPR binding by gp68 shown as sticks. **j**, Binding pocket of 1′′,3′-gcADPR in RyR Tad is shown. Polar interactions are colored as yellow dashed lines. **k**, Overlay of sensorgrams from SPR experiments used to determine the kinetics of RyR Tad and its mutants binding to 1′′,3′-gcADPR. Data were fitted with a model describing one-site binding for the ligands (black lines). **l**, Spot titration plaque assay of F10 phage on bacterial lawns with and without type I Thoeris defense, overexpressing RyR Tad wild type (WT) and its mutant version W132A. C-terminal domain of RyR Tad was used (gp68_C). Phage was titrated in 10-fold serial dilutions. **m**, Summary of the binding results of gp68. The detailed binding curves are shown in Extended Data Fig. 18.

We hypothesized that the RyR Tad C-terminal domain might antagonize type I Thoeris through binding to 1′′–3′ gcADPR. HPLC (High Performance Liquid Chromatography) confirmed binding of 1′′–3′ gcADPR by both full-length and the C-terminal domain of RyR Tad. Following inclusion of SDS, the bound molecule is released back into the buffer (Fig. 5f). To investigate the mechanism of sequestering 1′′–3′ gcADPR, we determined the crystal structure of RyR Tad bound with 1′′–3′ gcADPR (Extended Data Table 1 and Fig. 5g). In the crystal structure, RyR Tad folds as a symmetrical dimer each with two domains (Fig. 5g). The dimeric state was also confirmed in solution using SEC-MALS assay (Extended Data Fig. 17). Dali search^47^ returned the Repeat12 domain of RyR proteins as the most similar entries to the C-terminal domain as predicted. Phage RyR Tad C-terminal domain and mammalian RyR3 R12 have similar structures with an RMSD of 3.0 Å over 115 Cα atoms (Fig. 5h). For the RyR Tad dimer, both domains are involved in dimerization of the protein. Specifically, the N-terminal domain folds into a domain-swapped conformation, with its β1 strand inserted into the other protomer, forming an intact five-stranded β sheet (Fig. 5g). The C-terminal domain folds into an all-α helical structure, with its α1 and α4 from each protomer interacting in a back-to-back manner, mediating the dimerization (Fig. 5g). The binding pocket of 1′′–3′ gcADPR overlaps with that of Repeat12 domain of RYR3 (Fig. 5h). The position of some residues involved in ligand binding are also conserved between the phage and mammalian proteins (Fig. 5i,j and Fig. 5c). Mutations in the binding residues H129A/W132A, H154A/K147A, K170A/D171A displayed binding K_D_ of around 100 times weaker than that of WT protein (K_D_ = 43 nM), as shown by SPR (surface plasmon resonance, Fig. 5k). Substitution of Trp132 to alanine (W132A) abolished RyR Tad anti-Thoeris type I activity in bacteria (Fig. 5l). Because reported sponge proteins bind different signal molecules^21,22^, we also investigated the binding spectrum of RyR Tad. Interestingly, SPR showed that RyR Tad binds multiple signals, with a K_D_ lower than 1 µM for 1′′–2′ gcADPR, and cADPR, in addition to 1′′–3′ gcADPR (Fig. 5m and Extended Data Fig. 18). In sum, these data reveal a phage molecular sponge identified in a phage anti-signaling hotspot that binds small molecule targets to inhibit innate signaling in bacteria that has striking structural and sequence similarity to a small molecule binding region of the human ryanodine receptor (Extended Data Fig. 19).

## Discussion

Signaling-based anti-phage defense systems such as CBASS and Thoeris have emerged as central components of bacterial innate immunity, yet the mechanisms by which these systems detect infection and the strategies phages use to evade them remain incompletely understood. Here, we show that closely related *Migulavirinae* phages encode both variable immune-activating determinants and a concentrated hotspot of anti-signaling proteins that together shape the outcome of host-virus interactions. Our findings suggest that successful escape from signaling immunity is not solely determined by the presence or absence of an inhibitor, but rather by the balance between the strength of immune activation and the collective activity of multiple protective phage genes.

A major finding of this work is the identification of side tail fiber proteins as determinants of type II Thoeris activation. Through phage hybridization and spontaneous escape analyses, we found that sequence variation within the knob domains of *Migulavirinae* side tail fibers strongly alters susceptibility to type II Thoeris. These findings contribute to a broader emerging view that bacterial immune systems can function analogously to pattern-recognition pathways that discriminate conserved viral features^48,49^. Moreover, this work demonstrates that the Ab09 side tail fiber is a weaker activator than the Lit1 equivalent, and therefore when Lit1 (which also possesses the anti-defense locus) acquires this region from Ab09 it becomes type II Thoeris resistant. However, the Ab09 tail fiber still has some Thoeris-activating ability as this phage relies on Bes1 and Bes2 to achieve full type II Thoeris evasion.

*Migulavirinae* phages encode a compact island containing multiple genes that reverse the effects of CBASS and Thoeris systems through different mechanisms. The combined activities of Acb4-Sequestin, Bes1, and Bes2 protect Ab09 against type I Thoeris, type II Thoeris, and type II-A CBASS, demonstrating a striking degree of redundancy and functional overlap. Unlike Acb4-Sequestin, which contains recognizable cyclic nucleotide sponge domains and directly binds signaling molecules, Bes1 and Bes2 do not appear to measurably bind or degrade gcADPR or 3′,3′-cGAMP in vitro, nor do they trigger Panoptes. Their mechanisms therefore likely differ fundamentally from currently characterized signal sequestration strategies. *bes1 and bes2* may contribute to phage development in a way that enables evasion of signaling systems, without being inhibitors *per se.* Bes1 contains a conserved RuvC-like fold, raising the possibility that it interacts with nucleic acids or signaling-associated complexes, whereas Bes2 lacks clear similarity to known protein families. Together, these findings suggest that phages have evolved additional, mechanistically distinct strategies to suppress nucleotide-based immunity beyond direct signal sequestration.

The anti-signaling hotspot identified here is broadly conserved among N4-like phages infecting diverse bacterial hosts, highlighting this genomic region as a recurrent platform for acquisition of immune antagonists. Notably, the locus contains a diverse collection of known and predicted inhibitors, including Tad2, Tad3, Acb4, Sequestin, and Bes1 and Bes2 homologs. This organization parallels the clustering of anti-CRISPR genes in mobile genetic elements^41,50–52^. This hotspot also enabled our identification of a gcADPR sponge structurally related to the mammalian ryanodine receptor revealing an unexpected connection between phage anti-defense proteins and eukaryotic signaling biology. The gp68 C-terminal domain shares both sequence and structural similarity with the RyR Repeat12 region, a putative nucleotide-binding domain implicated drug and ATP binding as well as proposed to be a metabolic sensor. Our work demonstrates that the phage RyR Tad fold binds tightly to NAD derivatives, which have a selective advantage in phage biology. More broadly, these findings present the idea that cyclic nucleotide-binding architectures are evolutionarily versatile scaffolds that can be repurposed across deeply divergent biological systems.

Together, our study reveals that closely related phages modulate signaling immunity through both immune activation tuning and multi-layered signal antagonism. These findings establish *Migulavirinae* and related N4-like phages as a rich reservoir of anti-signaling mechanisms and provide new insight into how phages evolve to manipulate bacterial innate immune pathways.

## Extended Data Figures

**Extended Data Figure 1.**
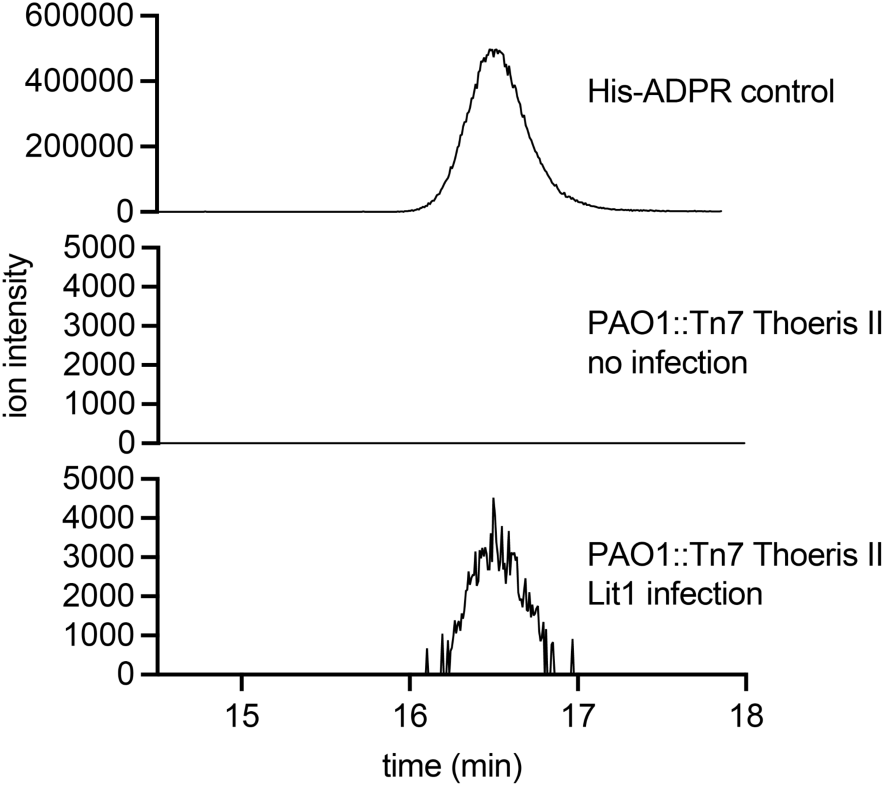
His-ADPR is detected in Lit1-infected *P. aeruginosa* harboring Thoeris II. Extracted ion chromatograms (EICs) of His-ADPR [m/z 695.1163–695.1303] in the lysate of *Bacillus subtilis* expressing ThsB from *Bacillus amyloliquefaciens* and infected with phage SPO1 (top), *P. aeruginosa* PAO1:Tn7 Thoeris II infected with no phage (middle), and *P. aeruginosa* PAO1:Tn7 Thoeris II infected with Lit1 phage (bottom).

**Extended Data Figure 2.**
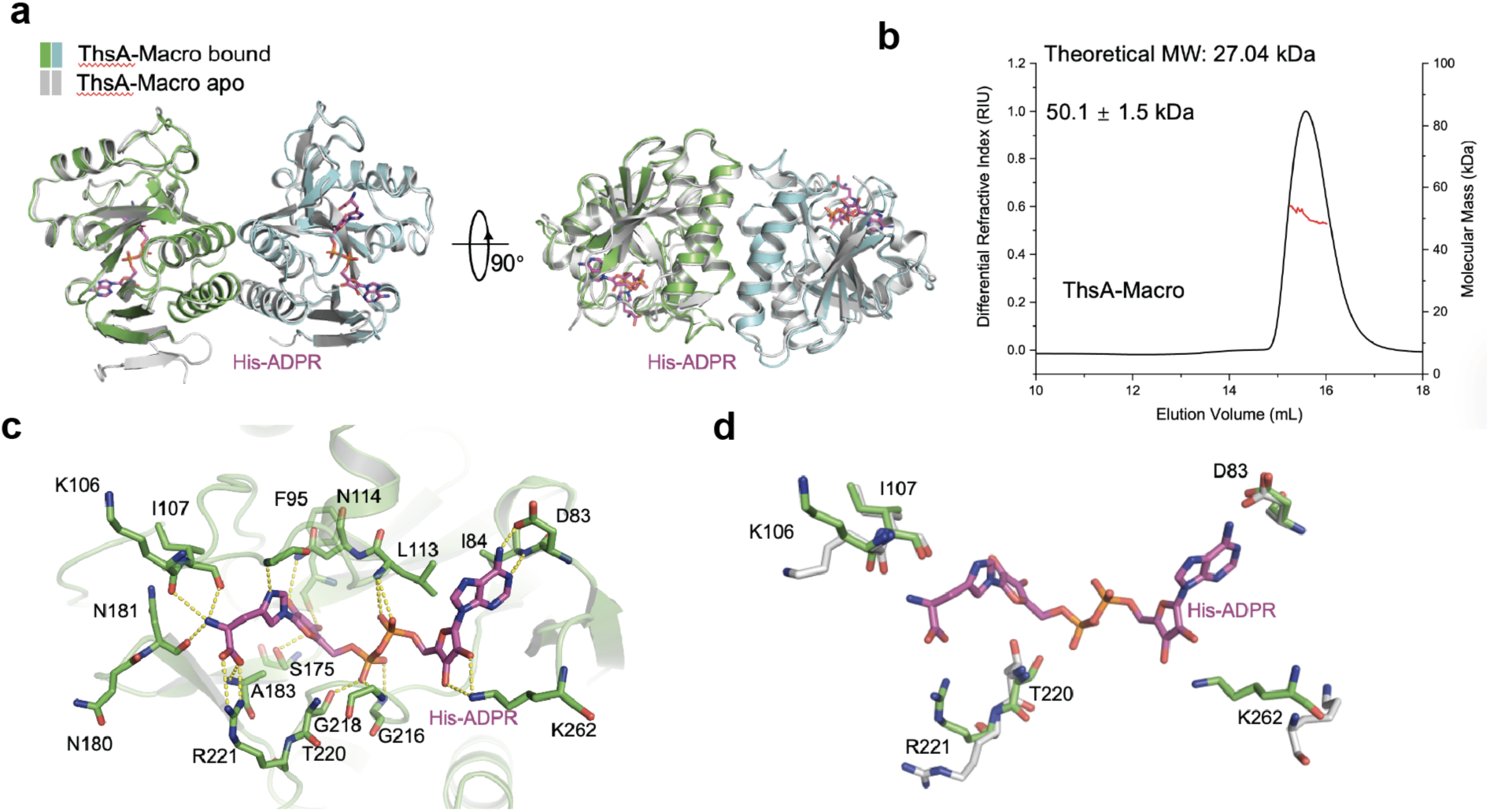
ThsA-Macro structure. **a**, Structural alignment of ThsA-Macro in the apo form and His-ADPR-bound form. His-ADPR is shown as sticks. **b**, SEC-MALS assay of purified ThsA-Macro. Calculated molecular weight of the sample and theoretical molecular weight of a monomer are shown. **c**, Binding pocket of His-ADPR in ThsA-Macro is shown. Polar interactions are colored as yellow dashed lines. **d**, Structural alignment of ThsA-Macro in the apo form and His-ADPR-bound form with residues displaying marked conformational changes between the two structures shown in sticks.

**Extended Data Figure 3.**
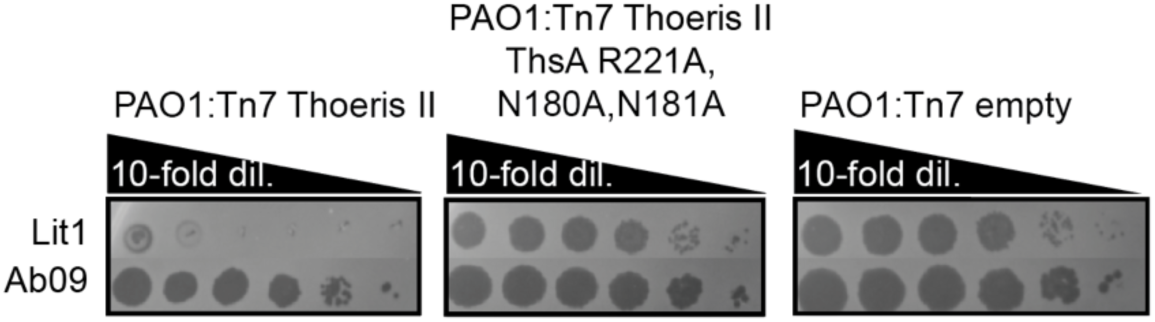
Mutations in ThsA active site abolish type II Thoeris defense. Spot titration plaque assay of indicated phages on bacterial lawns over-expressing the wild type type II Thoeris locus (PAO1:Tn7 Thoeris II). Indicated point mutations in ThsA-macro His-ADPR binding site (PAO1:Tn7 Thoeris II ThsA R221A, N180A, N181A), and with empty Tn7 vector integration in PAO1 genome (PAO1:Tn7 empty). Phages were titrated in 10-fold serial dilutions.

**Extended Data Figure 4.**
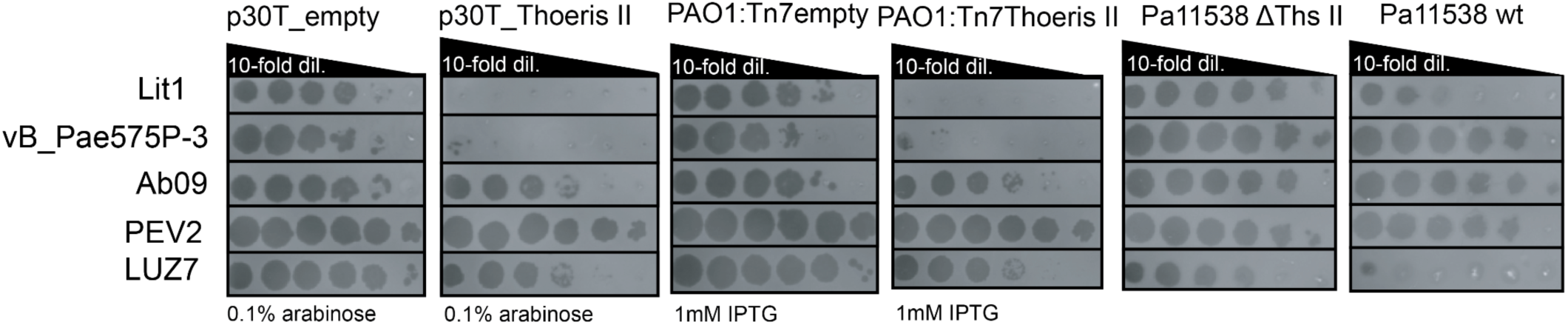
*Migulavirinae* bacteriophages are targeted with different efficiency in different Pa strains. POA1:Tn7 Thoeris II strain that over-express type II Thoeris MRSN11538 locus under regulation of IPTG-inducible promoter, PAO1 strain transformed with pHerd30T plasmid that encodes type II Thoeris MRSN11538 locus under the regulation of IPTG inducible promoter, and MRSN11538 native strain that naturally encodes type II Thoeris. Phages were titrated in 10-fold serial dilutions.

**Extended Data Figure 5.**
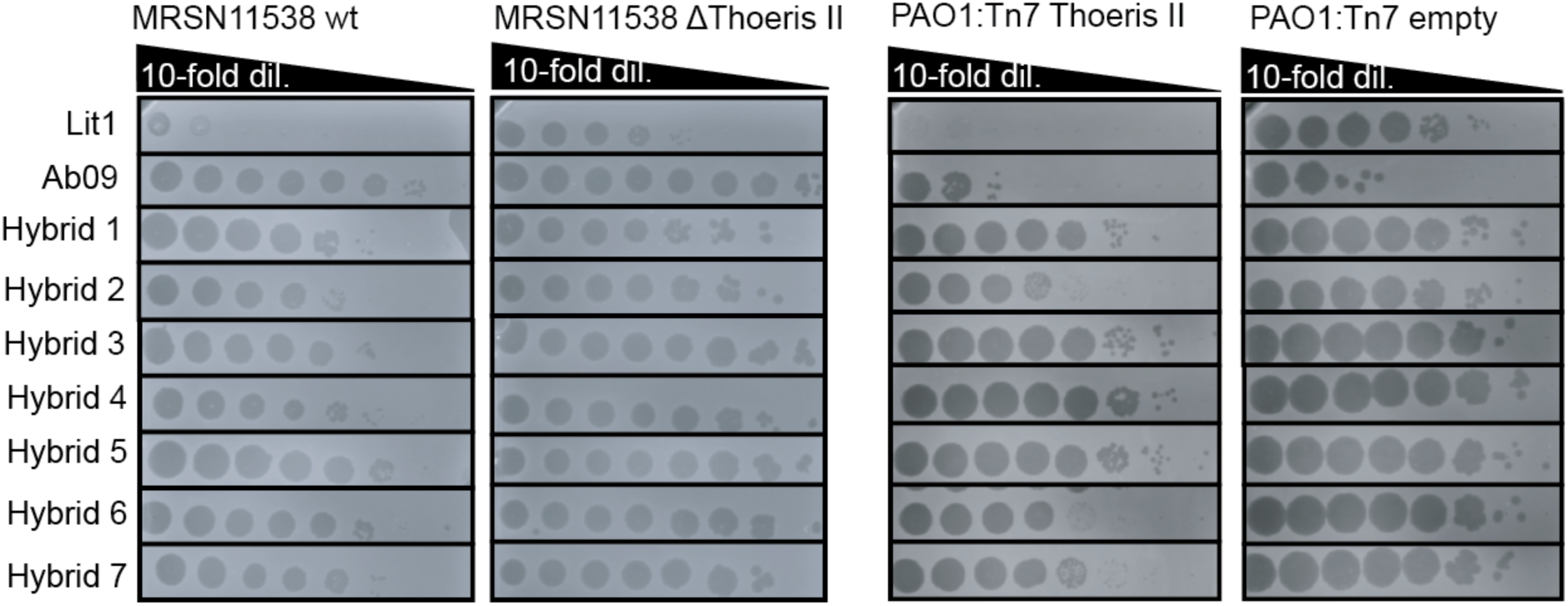
Resistance of Lit1/Ab09 hybrids to type II Thoeris in clinical strain MRSN11538 and in PAO1:Tn7 Thoeris II. Spot titration plaque assay of indicated phages on bacterial lawns with or without type II Thoeris: wild type MRSN11538 or MRSN11538 Δtype II Thoeris, PAO1 with type II Thoeris locus integrated in the genome (PAO1:Tn7 Thoeris II) or PAO1 without the defense (PAO1:Tn7 empty). Lit1/Ab09 hybrids sequences are shown in Fig. 2b. Phage were titrated in 10-fold serial dilutions.

**Extended Data Figure 6.**
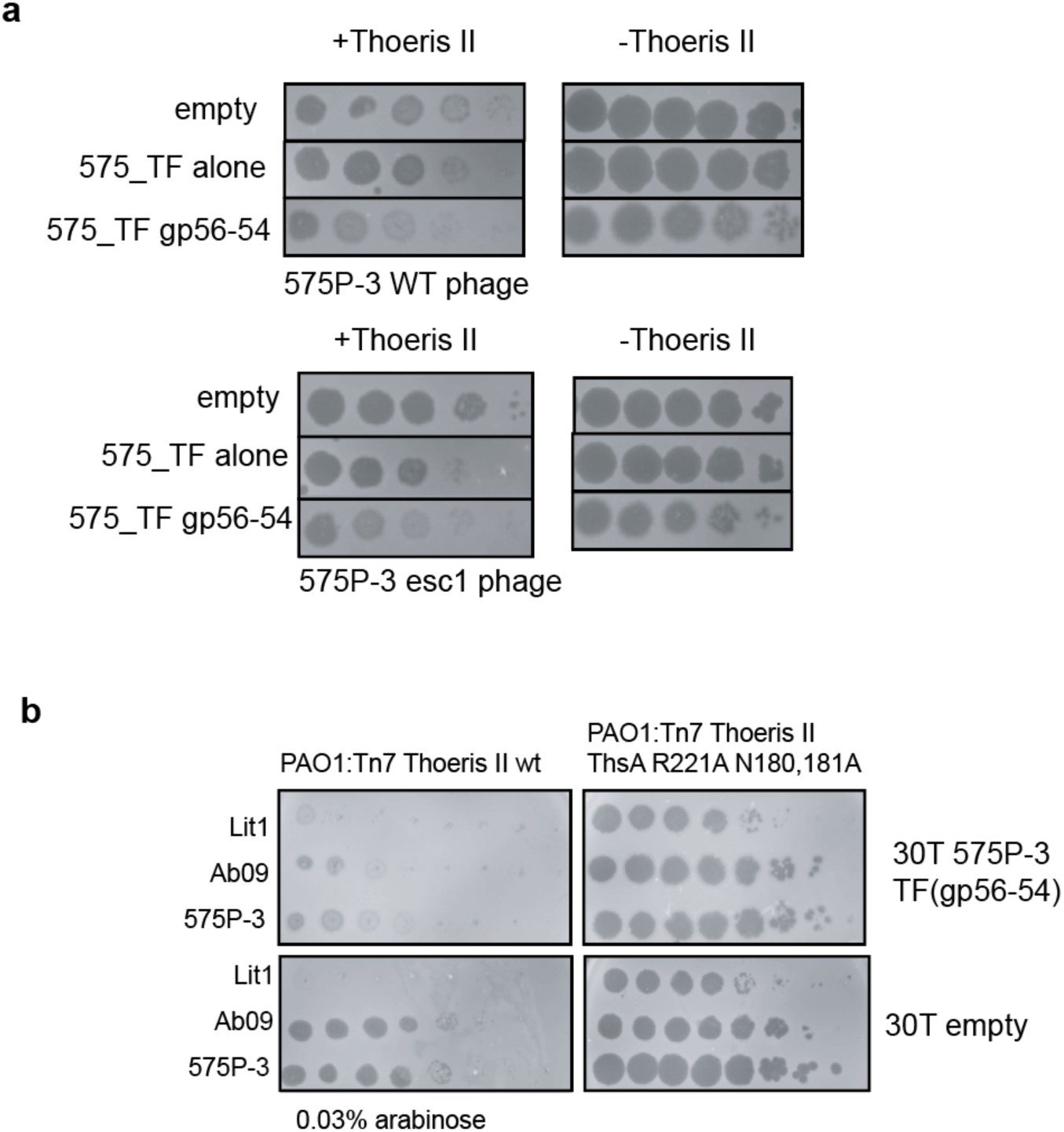
Activation of type II Thoeris through overexpression of the side tail fiber gene. **a**, Overexpression of 575P-3 side tail fiber alone or with downstream genes in the presence or absence of type II Thoeris. Spot titration plaque assay of indicated phages on bacterial lawns with wild type II Thoeris locus integrated in the genome (PAO1:Tn7 Thoeris II wt). **b**, Spot titration of the indicated phages on lawns expressing WT or mutant type II Thoeris and/or a tail tiber construct. Phage were titrated in 10-fold serial dilutions.

**Extended Data Figure 7.**
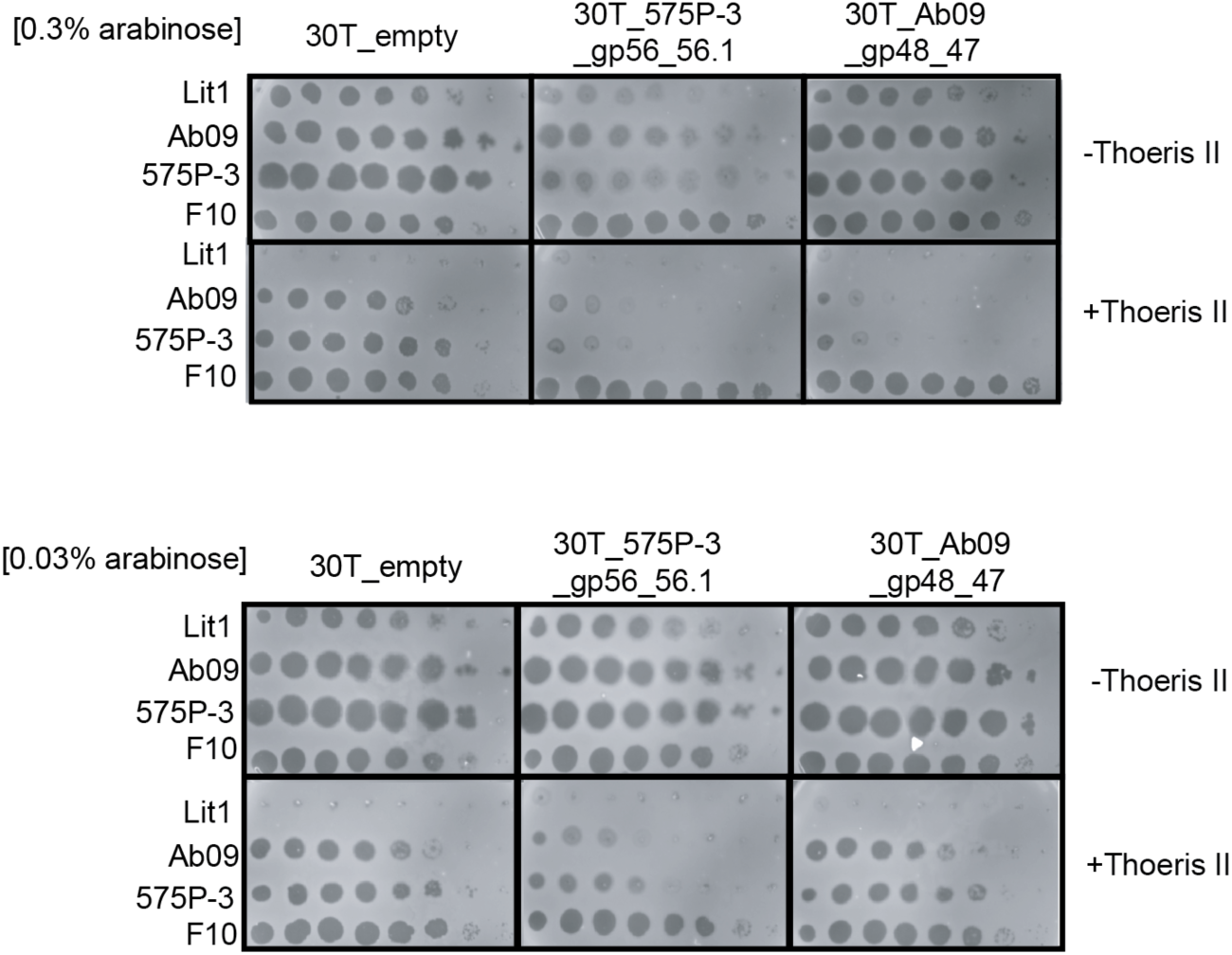
575P-3 side tail fiber with its chaperone activates type II Thoeris more efficiently than the corresponding genes from Ab09. Spot titration plaque assay of indicated phages on bacterial lawns induced with 0.3 mM IPTG, expressing the type II Thoeris locus integrated in the genome (Tn7 Thoeris II) or without (Tn7 empty). Indicated phage genes are over-expressed from 30T plasmids with 0.3% (top panels) or 0.03% arabinose (bottom panels). Phage were titrated in 10-fold serial dilutions. Arabinose includes the 30T construct while IPTG induces the Tn7 Thoeris system.

**Extended Data Figure 8.**
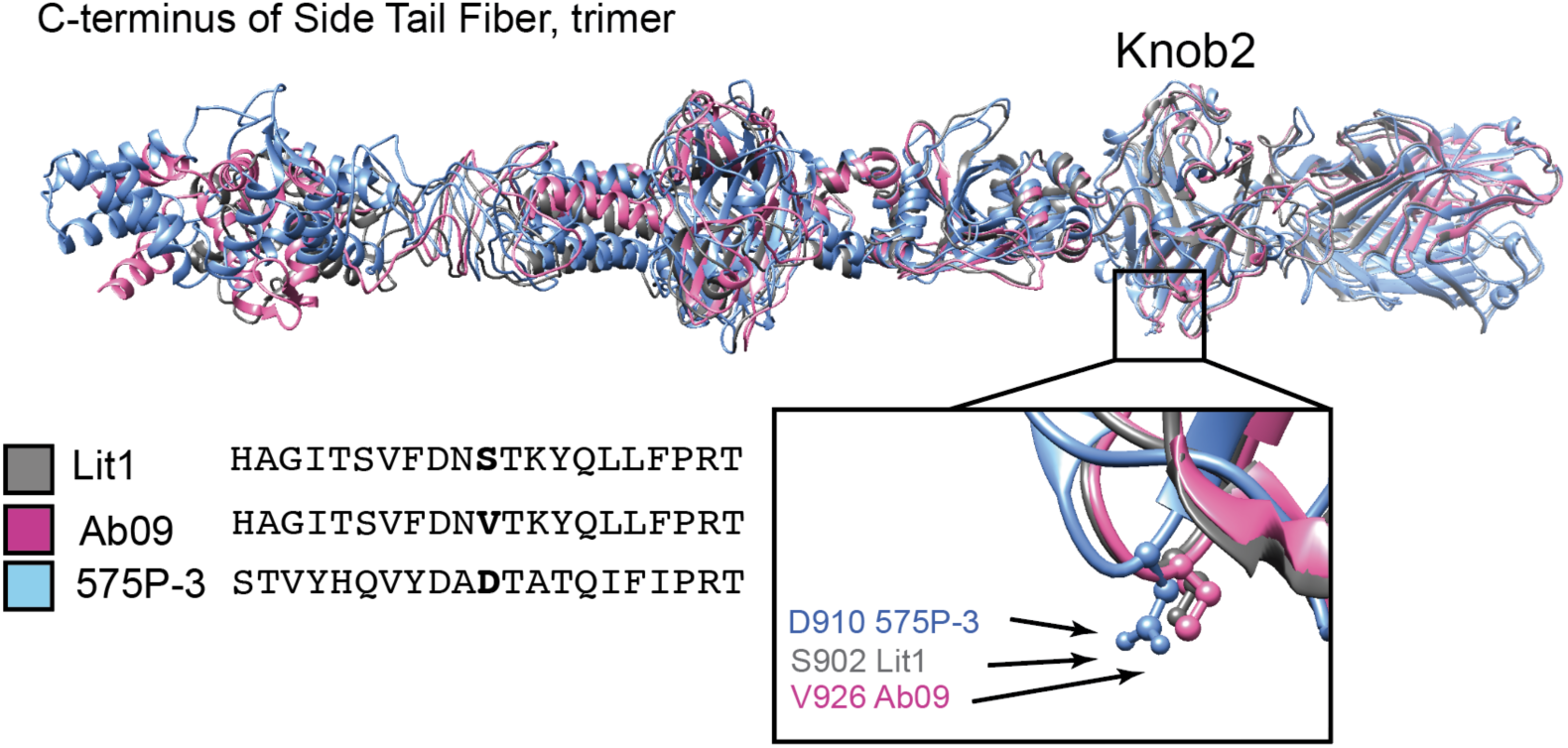
Structural alignment of C-terminal parts of Ab09, Lit1, 575P-3 side tail fibers. Lit1 is shown in grey, Ab09 in pink, 575P-3 in blue, position of 575P-3 D910 aspartic acid that allows type II Thoeris escape, as well as corresponding amino acids in Ab09 and Lit1 knob2 domains, are shown in the black rectangle. Amino acid sequences adjacent to the designated positions are shown.

**Extended Data Figure 9.**
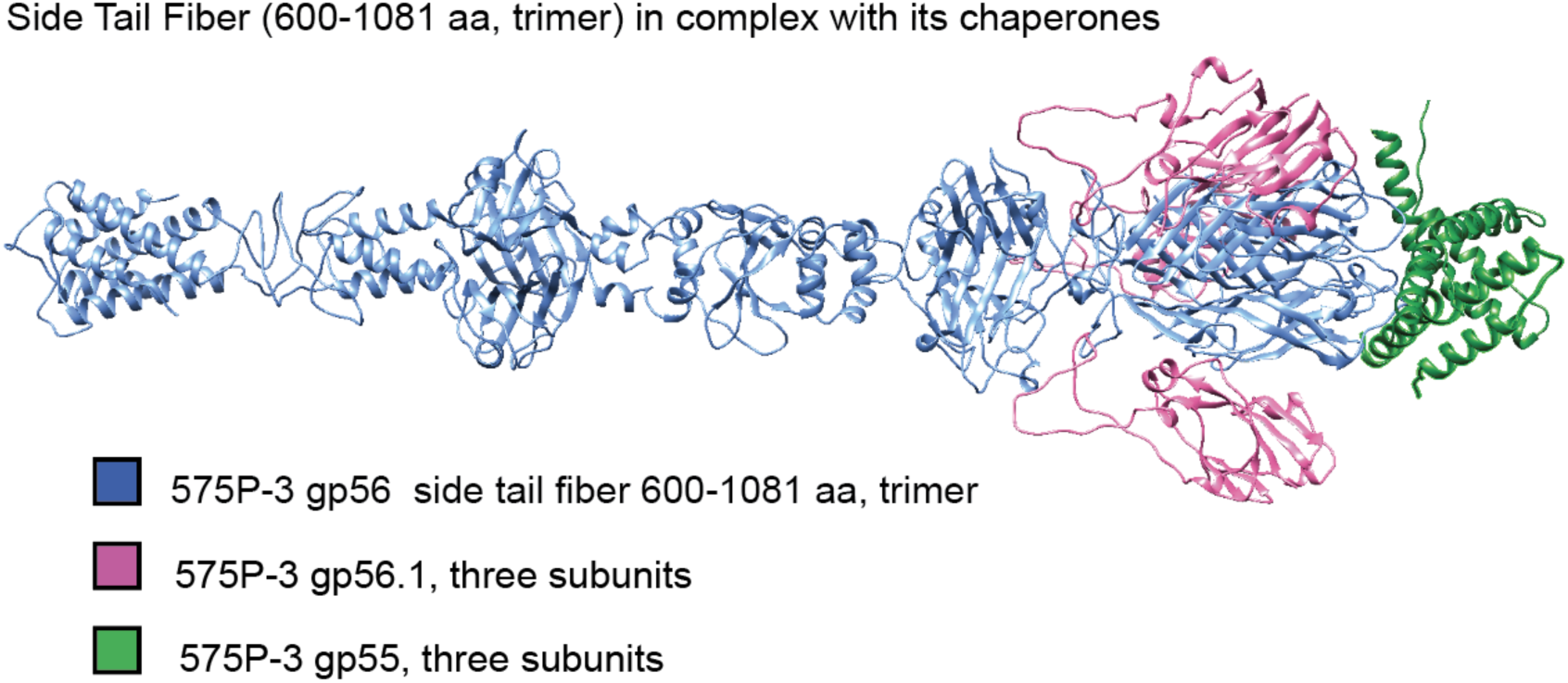
Ab09 side tail fiber gp48 forms a complex with gp47 and gp46 chaperones. AlphaFold model of gp48 trimer with three subunits of gp47and gp46.

**Extended Data Figure 10.**
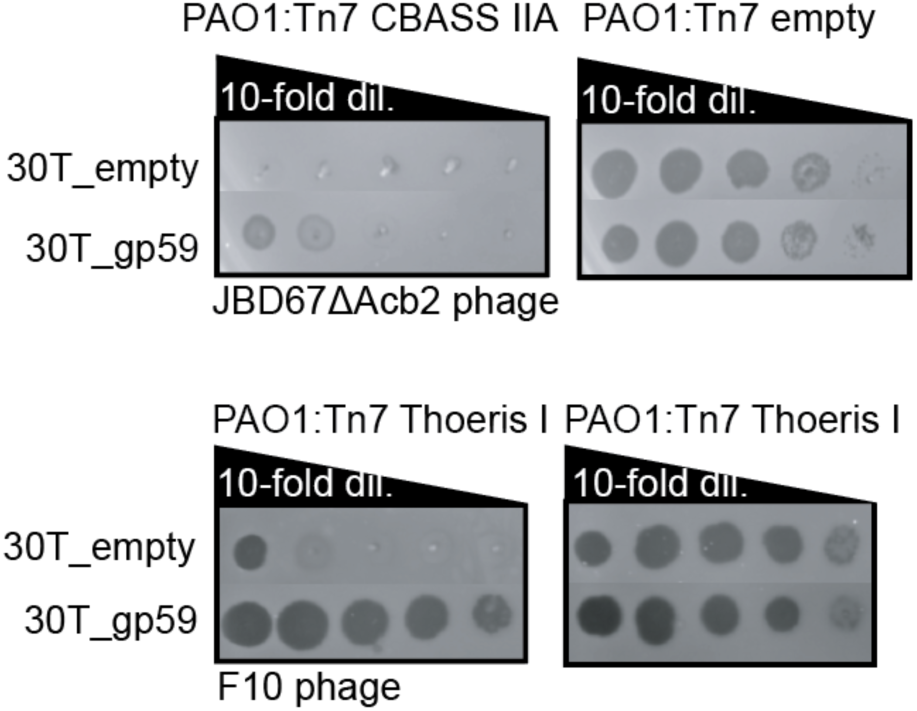
Ab09 gp59 protein, Acb4-Sequestin fusion, possesses modest anti-CBASS II-A and strong anti-Thoeris II activity. Spot titration plaque assay of indicated phages on bacterial lawns. Phages were titrated in 10-fold serial dilutions.

**Extended Data Figure 11.**
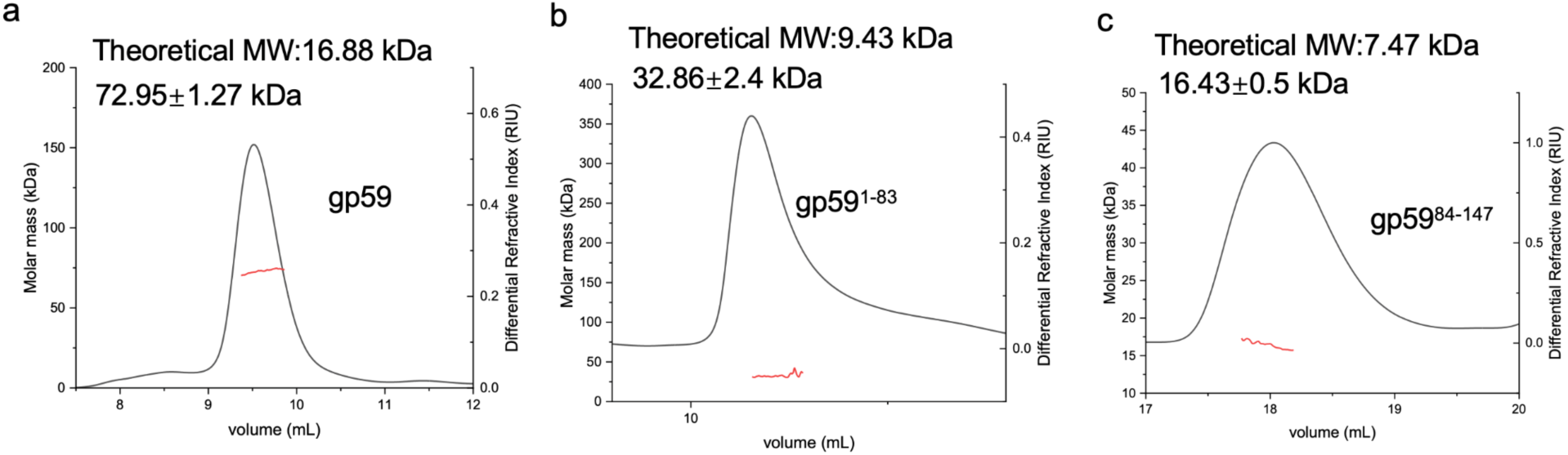
Oligomeric state of gp59 and its domains. SEC-MALS assays of purified gp59 and its two domains (Acb4 - gp59 1-83 and Sequestin gp59 84-147). Calculated molecular weight of the sample and theoretical molecular weight of a monomer are shown.

**Extended Data Figure 12.**
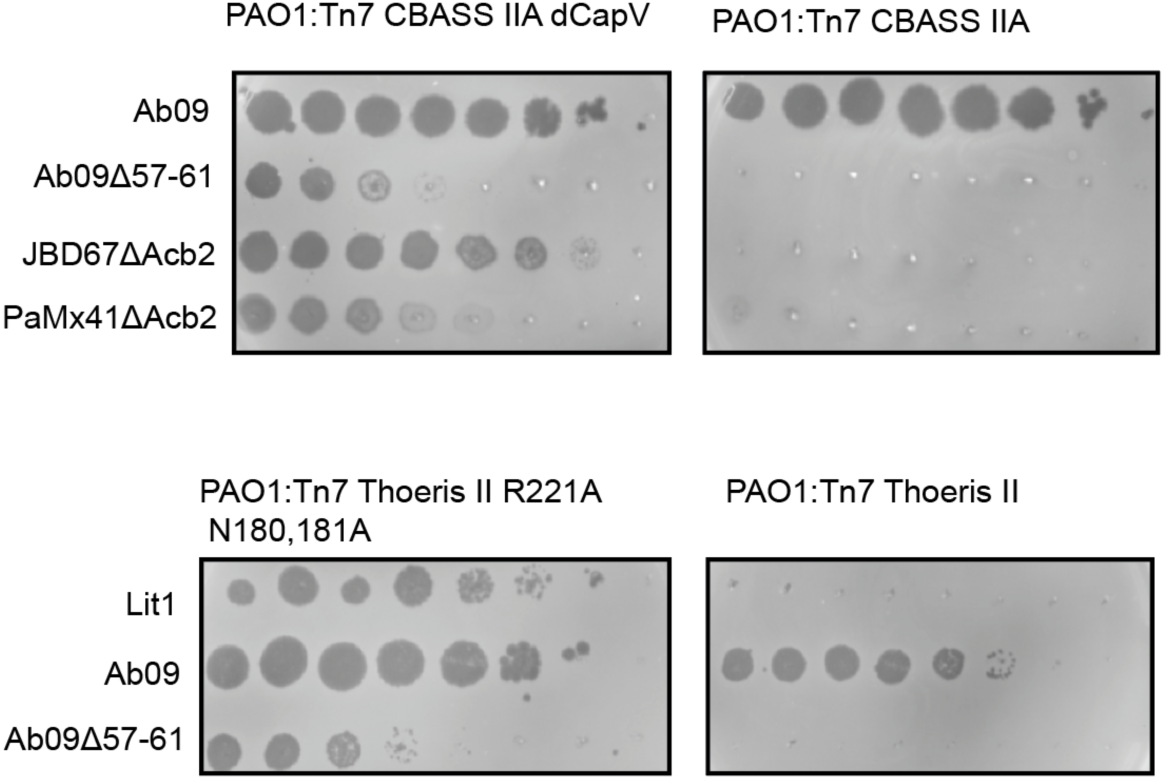
Targeting of Ab09Δ57-61 by type II Thoeris and type II-A CBASS is dependent on the presence of active effector proteins. Spot titration plaque assay of phages on bacterial lawns. Phage was titrated in 10-fold serial dilutions.

**Extended Data Figure 13.**
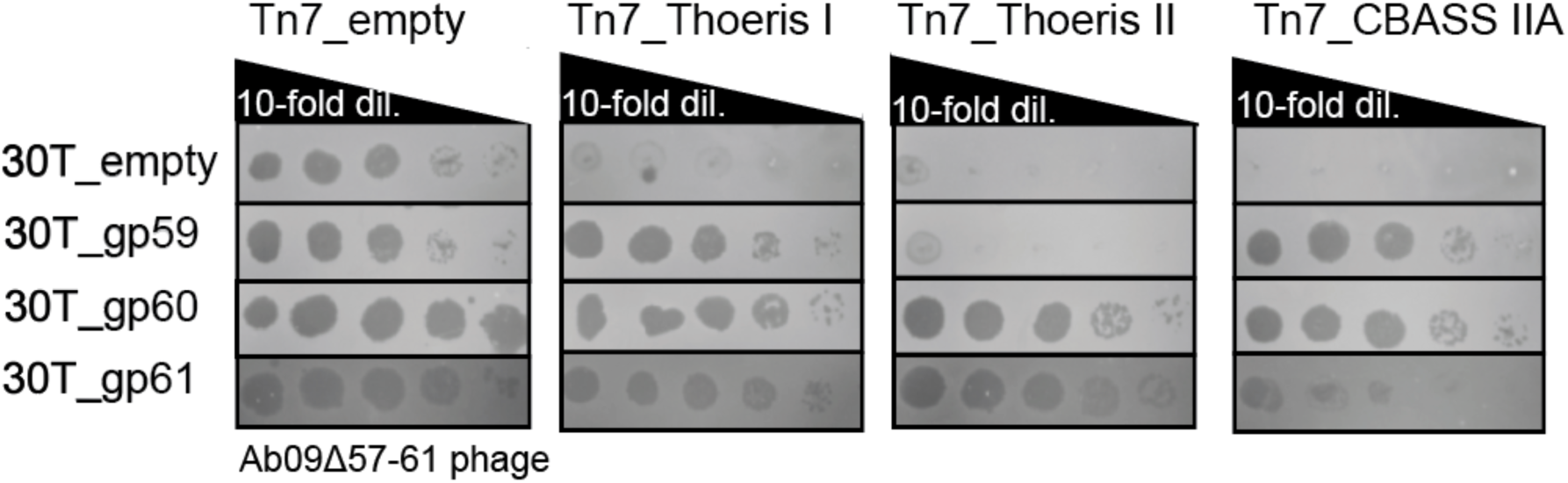
gp59,60 and 61 plasmid expression to assess complementation of phage Ab09*Δ57-61*. Each plasmid (30T) expressing the indicated gene has been used to transform strains with the indicated chromosomally expressed phage defense system (Tn7). Spot titration plaque assay of phages on bacterial lawns. Phage was titrated in 10-fold serial dilutions.

**Extended Data Figure 14.**
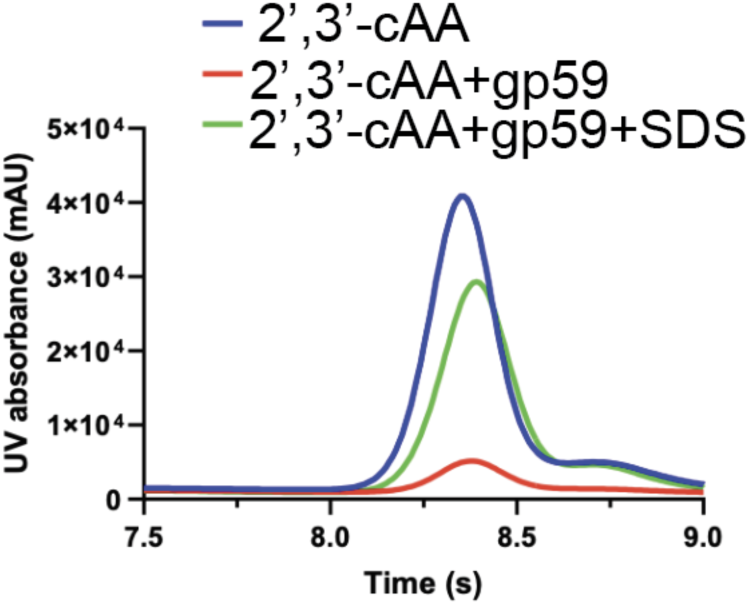
The ability of gp59 to bind and release 2′,3′-cAA when treated with SDS was analyzed using HPLC. 2′,3′-cAA standard was used as a control. The remaining nucleotides after incubation with gp59 were tested.

**Extended Data Figure 15.**
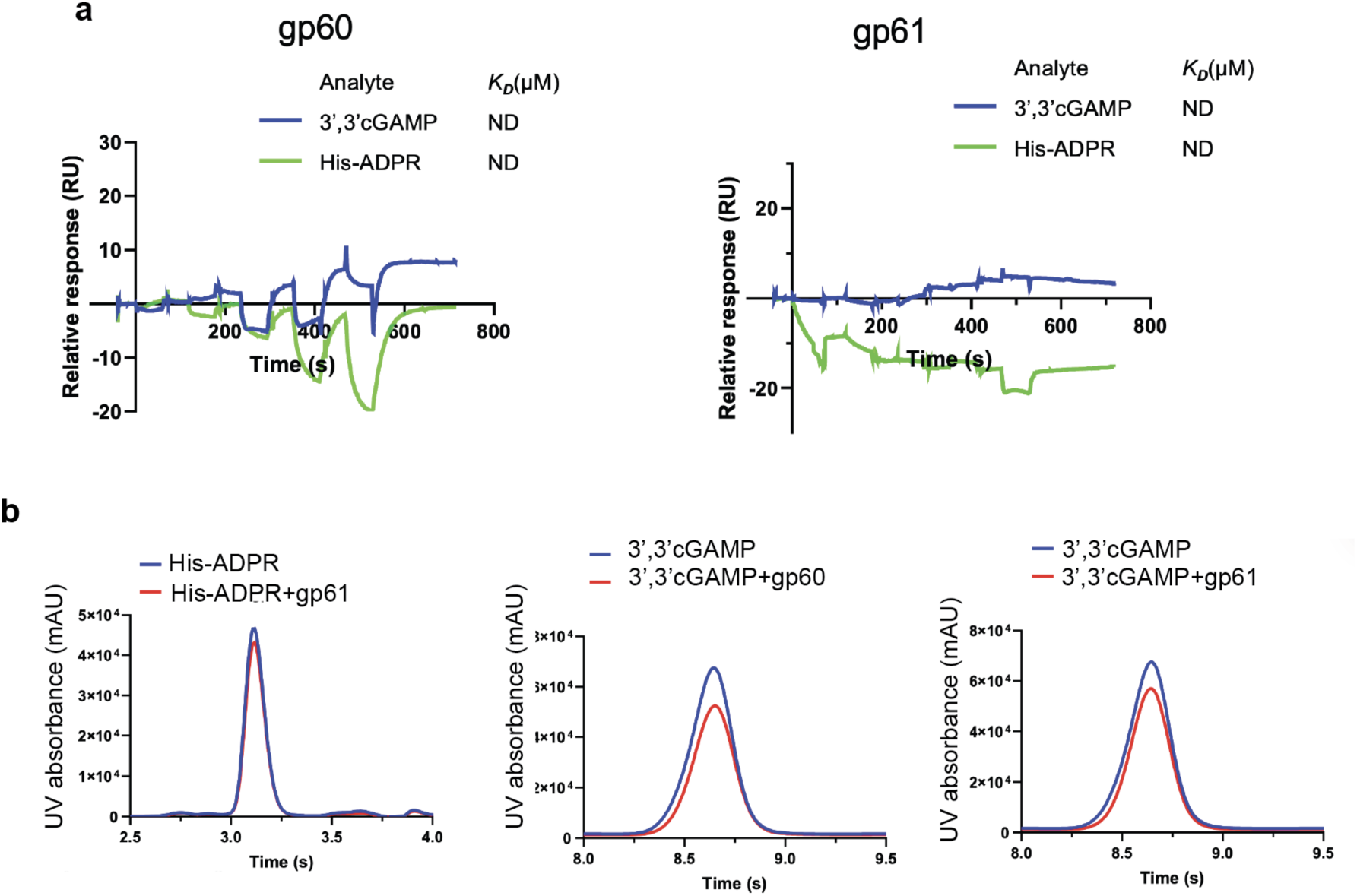
Bes1 and Bes2 do not bind type II Thoeris and type II-A CBASS signals *in vitro*. a, Overlay of sensorgrams from SPR experiments used to determine kinetics of gp60 (Bes1) and gp61 (Bes2) binding to 3′,3′cGAMP and His-ADPR. b, The ability of gp60 (Bes1) and gp61 (Bes2) to bind or cleave 3′,3′cGAMP and His-ADPR was analyzed using HPLC. The remaining nucleotides after incubation with gp60 or gp61 were tested.

**Extended Data Figure 16.**
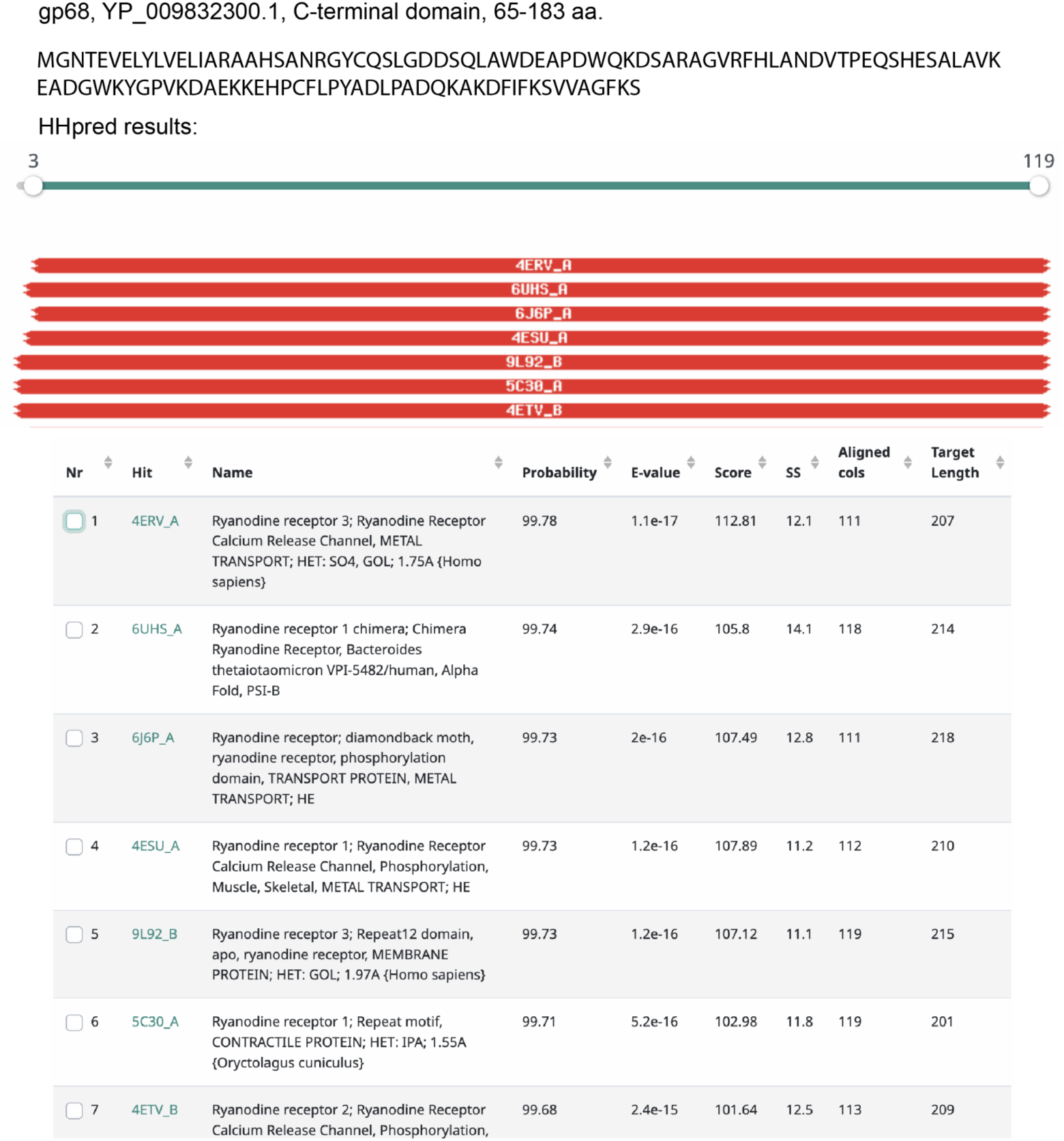
HHpred results showing the C terminal domain of gp68 is related to ryanodine receptors Repeat 12 domain.

**Extended Data Figure 17.**
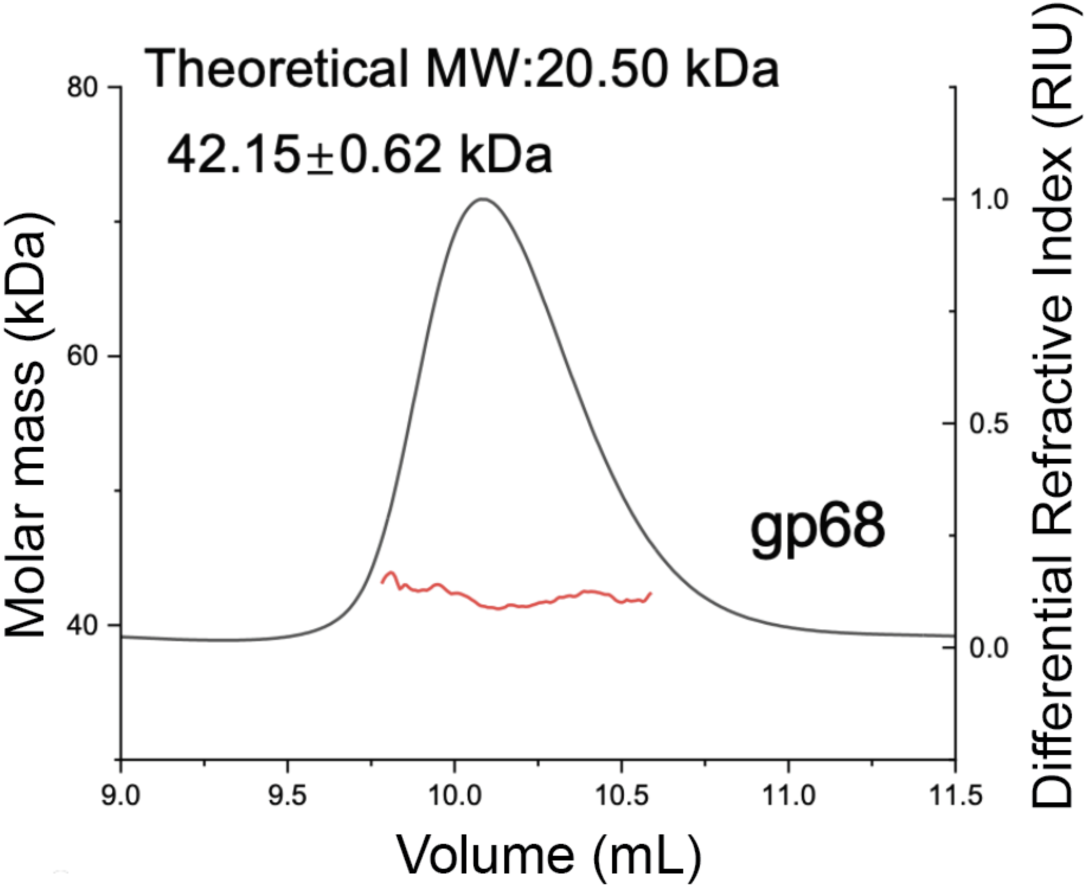
RyR Tad forms dimers in solution. SEC-MALS assay of purified gp68 (RyR Tad). Calculated molecular weight of the sample and theoretical molecular weight of a monomer are shown.

**Extended Data Figure 18.**
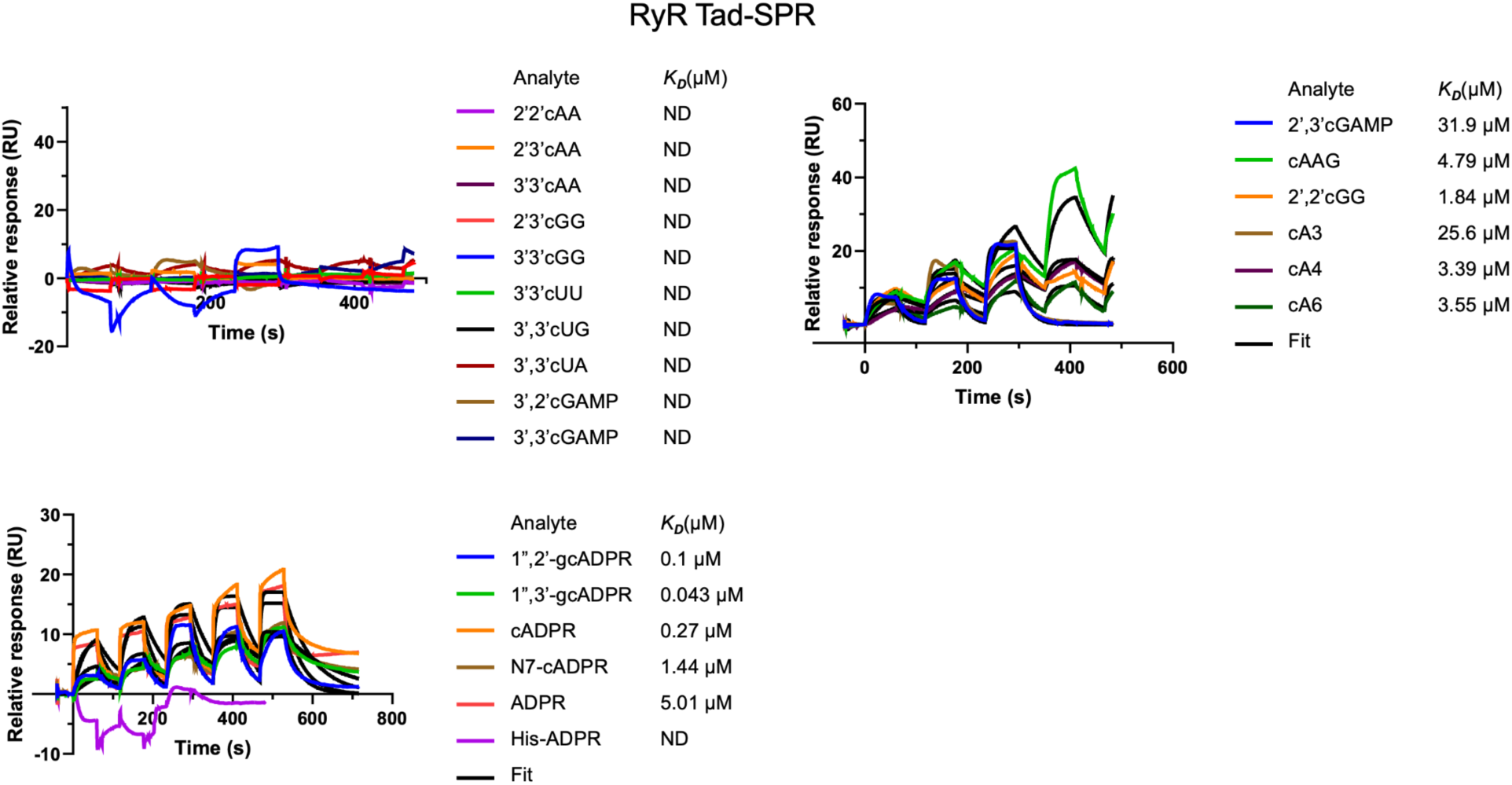
Overlay of sensorgrams from SPR experiments used to determine the kinetics of gp68 binding to various nucleotides. Data were fitted with a model describing one-site binding for the ligands (black lines).

**Extended Data Figure 19.**
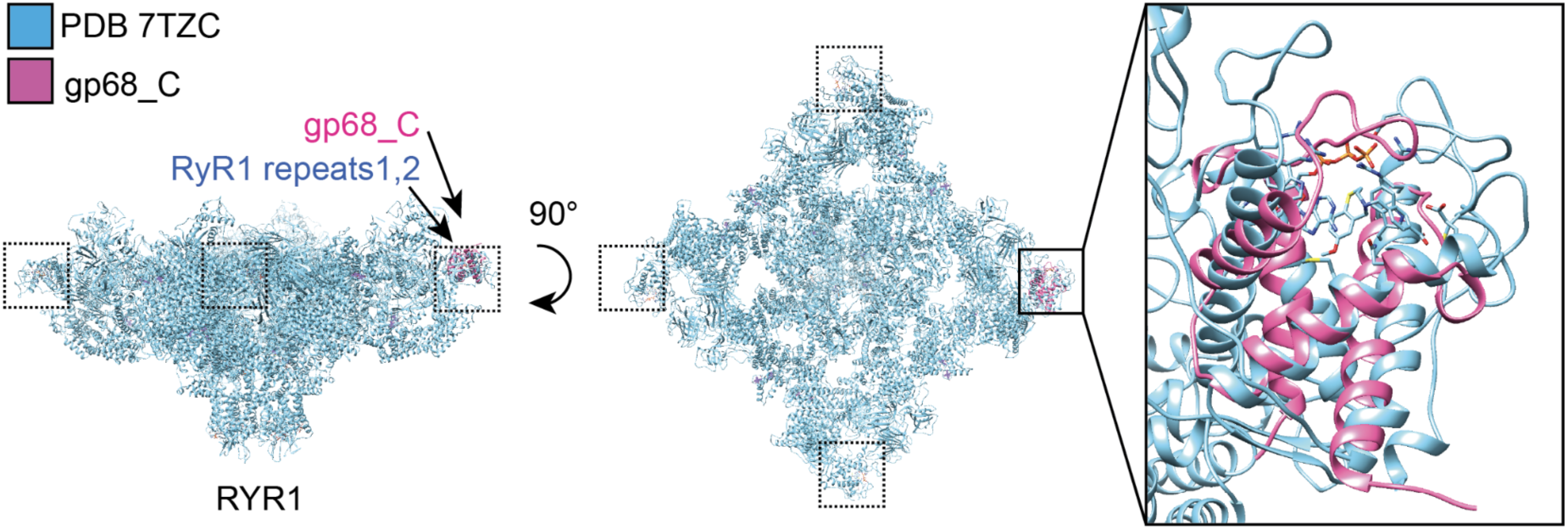
C-terminal domain of gp68 is structurally similar to RYR1 Repeat12 domain. Structural alignment of PDB 7TZC RYR1 (blue) with AlphaFold model of gp68_C terminal domain (pink) is shown. In box, RyR1 Repeat1,2 domain in complex with ATP aligned with gp68_C.

## Acknowledgements

We thank the ID02-Microfocusing X-Ray Protein Crystallography Beamline of High Energy Photon Source (https://cstr.cn/31138.02.HEPS.ID02) and beamlines BL02U1 and BL19U1 of the Shanghai Synchrotron Radiation Facility for providing technical support and assistance in data collection; and Y. Chen, Z. Yang and B. Zhou at the Institute of Biophysics, Chinese Academy of Sciences for technical help with SPR experiments. We thank Agata Jurczak-Kurek for providing vB-Pae575P-3, Christine Pourcel for Ab09 and Rob Lavigne for PEV2 for bacteriophages, Walter Reed Army Institute of Research for MRSN11538 strain, Petr Leiman for insights on co-folding of side tail fiber with its chaperone.

J.B.D. is supported by the US National Institutes of Health (R21AI168811 previously supported CBASS studies, R01AI167412 for phage host range determinants), and the Kleberg Foundation. J.B.D. also acknowledges generous research gifts and awards from the University of California, San Francisco: the Bowes Biomedical Investigator Award and the University of Sanghvi-Agarwal Innovation Award research gifts. Y.F. is supported by National key research and development program of China (2022YFC3401500 and 2022YFC2104800), Scientific Research Innovation Capability Support Project for Young Faculty (ZYGXQNJSKYCXNLZCXM-B1) and the Young Beijing Scholars Program.

J.P.G. acknowledges support from the US National Science Foundation (IOS-2143636).

## Author contributions

I.F. and J.B.-D. conceived of the project and supervised it along with Y.F.

I.F, Y.F. and J.B.-D. designed experiments.

I.F. performed microbiology, genetics, strain engineering, bioinformatics, and phage experiments Y.Y., Z.G., J.L. and X.Y. purified the proteins, grew and optimized the crystals, collected the diffraction data and performed biochemical assays. Y.F. solved the crystal structures with the help of H.W.

M.G. performed escape bacteriophage isolation and strain engineering

Z.Z. performed LC-MS experiments

D.D.N. assisted with in vivo phage experiments

J.G. supervised LC-MS assays.

I.F, Y.F., and J.B.-D. wrote the original manuscript.

I.F, Y.F., J.G. and J.B.-D. revised the manuscript.

## Methods

### Bacterial strains

The P. aeruginosa strains (MRSN11538, PAO1) and E. coli strains (XL1-Blue) were grown in Lysogeny broth (LB) medium at 37°C both with aeration at 225 r.p.m. Bacteria plating was performed on LB broth supplemented with gentamicin for maintaining pHERD30T plasmid (50 μg ml−1 for P. aeruginosa and 20 μg ml−1 for E. coli), as well as with 10 mM MgSO4 for phage spot assays. Gene expression in P. aeruginosa was induced by the addition of 0.1–0.3% L-arabinose or 0.2-1 mM isopropyl-β-D-thiogalactopyranoside IPTG unless stated otherwise. The E. coli BL21 (DE3) strain was used for recombinant protein overexpression and grown in Lysogeny broth (LB) medium. The cells were grown at 37°C until OD600nm reached 0.8 and then induced at 18°C for 12 h. Strains used in the study are listed in Extended Data Table 2.

### Episomal gene expression

The shuttle vector pHERD30T that replicates in P. aeruginosa and E. coli was used for episomal expression of proteins in P. aeruginosa strains. pHERD30T has an arabinose-inducible promoter and a selectable gentamicin marker. The vector was linearized using around-the-world PCR (in positions of NcoI and HindIII sites), treated with DpnI, and then purified. The inserts were amplified using phage stock or overnight bacterial culture as a DNA template and joined with a linearized pHERD30T vector using NEBuilder HiFi DNA Assembly (NEB) following the manufacturer’s protocol. The resulting plasmids were used to transform E. coli XL1-Blue cells. All plasmid constructs were verified by whole plasmid sequencing. *P.aeruginosa* PAO1 cells were electroporated with the corresponding vectors and used in the experiments.

### Chromosomal type II Thoeris integration, wild type and ThsA mutant version

For chromosomal insertion of the wild type MRSN11538 Thoeris II operon and its version with mutations in ThsA His-ADPR binding site at the Tn7 locus in P. aeruginosa PAO1(PAO1:Tn7 Thoeris II), the integrating vector pUC18-mini-Tn7T-LAC^53^ carrying Thoeris operon (wild type or mutant) and the transposase expressing helper plasmid pTNS3 were used. pUC18-mini-Tn7T-LAC empty vector was used for the creation of the control strain (PAO1: Tn7 empty). The vector was linearized using around-the-world PCR (in positions of HindIII and BamHI sites), treated with DpnI, and then purified. The insert was amplified using MRSN11538 overnight culture as a DNA template and joined with a linearized pUC18-mini-Tn7T-LAC vector using HiFi DNA Assembly (NEB) following the manufacturer’s protocol. The resulting plasmids were used to transform E. coli XL1-Blue cells. All plasmid constructs were verified by whole plasmid sequencing. To introduce point mutations into the ThsA sequence, the pUC18-mini-Tn7T-LAC vector carrying the Thoeris II operon was linearized by around-the-world PCR using primers containing the desired substitutions at the corresponding positions. The N180A and N181A mutations were introduced first, followed by R221A in a second round of mutagenesis. P.aeruginosa PAO1 cells were electroporated with pUC18-mini-Tn7T-LAC and pTNS3 and selected on gentamicin-containing plates. Potential integrants were screened by colony PCR with primers PTn7R and PglmS-down^53^. Electrocompetent cell preparations, transformations, integrations, selections, plasmid curing, and FLP-recombinase-mediated marker excision with pFLP were performed as described previously^53^.

### Phage growth

Phages Lit1, Ab09, 575P-3, LUZ7, PEV2, F10 were grown on P. aeruginosa PAO1, which lacks CBASS and Thoeris systems. For experiments in P. aeruginosa MRSN11538 strain bacteriophages were propagated in MRSN11538 with deletion of type Thoeris operon. For phage propagation 100 µl of P. aeruginosa overnight cultures were infected with 10 µl of low titer phage lysate and then mixed with 3 ml of 0.35% top agar 10 mM MgSO_4_ for plating on the LB solid agar (20 ml LB agar with 10 mM MgSO_4_). After incubating 37 °C overnight, 2.5 ml SM phage buffer was added on the solid agar lawn and then incubated for 10 minutes at room temperature. The whole cell lysate was collected, a 10% volume of chloroform was added, and the tubes were left for 20 minutes at room temperature with gentle shaking, followed by centrifugation at maximum speed for 3 min 4°C to remove cell debris. The supernatant phage lysate was stored at 4°C for downstream assays.

### Plaque assays

Plaque assays were conducted at 37 °C with solid LB agar plates supplemented with 10 mM MgSO_4_, 50 µg ml-1 gentamicin, 0.1% L-arabinose (for gene expression from pHerd30T), and 0.3 mM IPTG for PAO1 strains with type I/ II Thoeris and type III-C CBASS operon chromosomal integration, and the same conditions except without IPTG for type II-A CBASS, unless otherwise indicated. Plaque assays in native MRSN11538 type II Thoeris strain were conducted at 37 °C with solid LB agar plates supplemented with 60 mM MgSO_4_ without IPTG. Note that type II Thoeris activity was impacted by MgSO_4_ levels with stronger targeting observed when 30-60mM MgSO_4_ is added to solid agar LB or liquid LB media. For induction of Panoptes genes 0.1% arabinose was used. 100 μL of overnight bacterial culture was mixed with top agar (0.35% agar in LB) and plated. Phage lysates were diluted 10-fold then 2 μL spots were applied to the top agar after it had been poured and solidified. The plates were incubated overnight at 37 °C.

### Lit1 and Ab09 phage hybridization assay

To generate Lit1 and Ab09 hybrid phages resistant to type II Thoeris both phages were used to infect MRSN11538 strain that naturally encodes Thoeris (MOI=1) and transformed with plasmid carrying type I-C CRISPR-Cas3 system targeting Ab09 *gp80* (absent in Lit1) protospacer sequence 5′-catgccctggtagaccccaaccatctctgctatg-3′ (based on Addgene 133773 pCas3cRh vector). Bacteriophages were added to 1 ml of MRSN11538 carrying pCas3cRh_target_gp80 OD=0.7 and incubated for 10 minutes at room temperature. Then 100 µl of infectious mix was mixed with 0.35% top agar with 60 mM MgSO_4_, plated on solid agar plate supplemented with 50 µg ml-1 gentamicin, 0.1% rhamnose (for expression of CRISPR-Cas genes). The plates were incubated overnight at 37 °C. The surviving bacteriophages were propagated to a high titer on the same selection strain. Genomic DNA was extracted from phage lysates using SDS/proteinase K treatment followed by column purification. For this 300 μL of high-titer phage lysate were mixed 1:1 with lysis buffer (final concentration: 10 mM Tris-HCl pH 8.0, 10 mM EDTA, 100 μg/mL proteinase K, 100 μg/mL RNase A, 0.5% SDS), incubated at 37 °C for 30 min and at 55 °C for 30 min, and then processed Genomic DNA purification Zymo kit. Sequencing was performed on an Illumina MiSeq instrument. Sequencing reads were aligned to Ab09 and Lit1 genomes using BWA aligner^54^. The position of Ab09 required regions was determined by visual analysis of hybrid sequence aligned to Ab09 and Lit1 phages in IGV Viewer^55^. Sequencing of Lit1 bacteriophage showed that phage that is used in this study has insertion of “G” in 30,885 position of genome preventing formation of premature stop codon in Lit1 side tail fiber gene in NCBI Lit1 genome file (NC_013692.1), resulting in full length of side tail fiber protein, similar to side tail fibers of other Migulavirinae members. The corrected sequence was used in all figures showing the Lit1 genome.

### Recombineering of Lit1 phage using an HDR template carrying the Ab09 side tail fiber genes

To introduce Ab09 gp48-45 sequences in Lit1 genome vector, p30T_HDR_Ab09_gp48_45 was cloned, which carries Ab09 gp48_45 genes introduced in the NcoI and HindIII sites of pHerd_30T vector. p30T_HDR_Ab09_gp48_45 was electroporated to PAO1 P. aeruginosa strain using 50 µg ml-1 gentamicin for selection. Lit1 bacteriophage was propagated on PAO1 p30T_HDR_Ab09_gp48_45 in the presence of 50 µg ml-1 gentamicin without the addition of inducers. The resulting phage lysate was used to infect the PAO1: Tn7_Thoeris II strain, with the addition of 0.3 mM IPTG to induce defense genes. The surviving Lit1 side tail fiber hybrids were isolated from single plaques and propagated to high titer using the same type II Thoeris overexpression strain. The genomic DNA was extracted, sequenced, and analyzed for the presence of Ab09-derived regions.

### Introducing deletions in phage

To introduce deletion without boundaries in Ab09 gp59 encoding region we used type I-C CRISPR-Cas system targeting gp59 (gp59_protospacer 5′-gatgctgagatcggcaaagaagtcgctcgtcgta-3′) with a mutation in Cas3 that abolishes helicase activity. Ab09 was propagated in PAO1:Tn7CRISPR-Cas3 that carries plasmid expressing crRNA targeting gp59_protospacer. Plaques resistant to CRISPR-Cas3 targeting, presumably due to loss of the protospacer region, were selected based on PCR analysis of the gp59-adjacent locus length. Deletion phages were sequenced and compared with the wild-type Ab09 genome to determine the boundaries of the deletions. The resulting phage is designated as Ab09*Δgp57-61.* For precise deletion of gp59, gp60–61, or gp59–61 from the Ab09 genome, HDR template plasmids p30T_HDR_gp59, p30T_HDR_gp60-61, and p30T_HDR_gp59-61 were generated, each carrying 600-bp homology arms flanking the corresponding target region. Ab09 was propagated on P. aeruginosa PAO1 carrying the corresponding HDR template plasmid to promote homologous recombination. The resulting phage lysates were then selected for deletion variants by propagation on P. aeruginosa PAO1 carrying a type I-C CRISPR-Cas system targeting the corresponding genes. Surviving bacteriophages were screened by PCR to assess the size of the deleted region, further propagated to high titer, and confirmed by whole-genome sequencing.

### Bioinformatic identification of hot spots of gene acquisition in N4-related phages

To identify putative anti-defense hotspots in N4-like bacteriophages, the online Gaia/SeqHub^43^ portal was used in co-search mode. The virion RNA polymerase gene (gp66 in Ab09) and the DNA primase-polymerase gene (gp55 in Ab09) were used as boundary markers. Genomes were selected for further analysis if they contained boundary genes with sequence and structural similarity to the Ab09 markers, encoded RecA and SSB-like genes adjacent to the DNA primase-polymerase gene, and had genome sizes of 60–80 kb. The identified hotspots were consistently associated with N4-related bacteriophages. ORFs encoded within the hotspot region were analyzed using Gaia/SeqHub, AlphaFold structural prediction, and Dali-based structural similarity searches. Genes encoding proteins encoded in the hot spot were considered candidate anti-signaling genes and were further tested in plaque assays for their ability to inhibit type I/II Thoeris or type II-A/III-C CBASS defense systems.

### LC-MS detection of His-ADPR

PAO1:Tn7 Thoeris II cells were grown in 50 ml of LB media supplemented with 0.5 mM IPTG and 10 mM MgSO_4_ until OD=0.6. Lit1 (a bacteriophage propagated on PAO1) was used to infect the cells (MOI=5), whereas no phage was added in the control sample without infection. Infected and control cells were incubated at 37 °C with shaking for 25 minutes and were centrifuged at 4 °C, 4,000 x g for 5 minutes. After the centrifugation, the supernatant was discarded and the pellet was resuspended in 600 µl of 100 mM Phosphate buffer (pH=7). Cells were sonicated in the presence of glass beads, and lysates were centrifuged at 4 °C , 16,000 x g for 20 minutes. 400 µl of the supernatant were transferred to Amicon Ultra-0.5 Centrifugal Filter Unit 3 kDa and centrifuged for 45 min at 4 °C, 12,000 x g. The samples were frozen at minus 80 °C. The liquid chromatography analysis was performed on a Thermo Vanquish UPLC using a Phenomenex Luna Omega 5 μm Polar C18 100 Å column (250×4.6 mm). The mobile phase A was water + 0.1 % (v/v) formic acid and the mobile phase B was acetonitrile + 0.1 % (v/v) formic acid. The flow rate was kept at 0.7 ml·min–1 and the gradient was as follows: 0% B (0–10 min), increase to 2.5% B (10–15 min), increase to 5% B (15–16 min), hold 5% B (16–26 min), increase to 95% B (26–27 min), hold 95% B (27–37 min), decrease to 0% B (37–38 min), hold 0% B (38–48 min). High-resolution electrospray ionization (HR-ESI) mass spectra were obtained using a Thermo Q Exactive Plus Hybrid Quadrople-Orbitap. The instrument was operated at negative ionization mode. The MS spectra were obtained on the orbitrap analyzer with a mass scanning range of 500–700 Da and resolution of 70,000, and then analyzed using mzmine 4.3.0 software. A cell lysate known to contain His-ADPR (*Bacillus subtilis* expressing ThsB from the *B. amyloliquefaciens* Y2 Thoeris system and infected with phage SPO1)^28^ was used as a standard.

### Protein expression and purification

In order to obtain the *Pae*ThsA^macro^ protein in apo state, the *PaethsA*^macro^ gene was cloned into a pET28a vector and carried an N-terminal 6×His tag. To obtain the *Pae*ThsA^macro^ protein bound to His-ADPR, we co-transformed an unlabeled *PaethsB* placed on the pET22b vector with it. To obtain the protein in the apo state, the genes of gp59 and its truncated forms, gp60, gp61, gp68 and its truncated forms were cloned into the pET28a vector with an N-terminal 6×His tag. All of the proteins were expressed in *E. coli* strain BL21 (DE3) and induced by 0.2 mM IPTG when the cell density reached an OD600 of 0.8. After growth at 18 °C for 12 h, the cells were collected, resuspended in lysis buffer (50 mM Tris-HCl pH 8.0, 300 mM NaCl, 10 mM imidazole and 1 mM PMSF) and lysed by sonication. The cell lysate was centrifuged at 20,000g for 50 min at 4°C to remove cell debris. The supernatant was applied onto a self-packaged Ni-affinity column (2 mL Ni-NTA, Genscript) and contaminant proteins were removed with wash buffer (50 mM Tris pH 8.0, 300 mM NaCl, 30 mM imidazole). The protein was then eluted with elute buffer (50 mM Tris pH 8.0, 300 mM NaCl, 300 mM imidazole). The eluant of protein was concentrated and further purified using a Superdex-200 increase 10/300 GL (GE Healthcare) column equilibrated with a buffer containing 10 mM Tris-HCl pH 8.0, 200 mM NaCl and 5 mM DTT. The purified proteins were analysed by SDS–PAGE. The fractions containing the target protein were pooled and concentrated.

### Crystallization, data collection and structural determination

After size-exclusion chromatography purification, the fractions containing the target protein were pooled and concentrated to 20 mg/mL. All crystals were obtained using the sitting-drop vapor diffusion method at 18°C by mixing 0.8 μL of protein solution with 0.8 μL of reservoir solution.

1. For both the apo form and His-ADPR-bound form of PaeThsAmacro, the reservoir solution contained 0.1 M Sodium chloride, 0.1 M Sodium formate, 0.1 M Bis-Tris propane pH 8.5 and 25% v/v PEG Smear Medium. Before harvesting, the crystals were cryoprotected in the reservoir solution supplemented with 20% (v/v) glycerol and then flash-frozen in liquid nitrogen.
2. For 1′′,3′-gcADPR-bound form gp68, the gp68 protein was mixed with 1′′,3′-gcADPR with a molar ratio of 1: 1.5 overnight before crystallization. The reservoir solution contained 0.2 M Sodium chloride, 0.1 M Bis-Tris pH 5.5 and 25% w/v PEG 3350. Before harvesting, the crystals were cryoprotected in the reservoir solution supplemented with 20% (v/v) glycerol and then flash-frozen in liquid nitrogen
3. For gp59 apo state, the reservoir solution contained 0.2 M Sodium chloride,0.05 M Bis-Tris pH 8.5 and 18% w/v PEG 3350. Before harvesting, the crystals were cryoprotected in the reservoir solution supplemented with 20% (v/v) glycerol and then flash-frozen in liquid nitrogen.

All the data were collected at the High Energy Photon Source (HEPS) in Beijing and beamline BL02U1 at the Shanghai Synchrotron Radiation Facility (SSRF), integrated and scaled using the HKL2000 package. The initial models of the three proteins were all obtained through modeling using AlphaFold3^56^. All the structures were solved through molecular replacement and refined manually using COOT^57^. The structure was further refined with PHENIX^58^ using non-crystallographic symmetry and stereochemistry information as restraints. The final structure was obtained through several rounds of refinement. Data collection and structure refinement statistics are summarized in Extended Data Table 1.

### SPR assay

The SPR analysis was performed using a Biacore 8K (GE Healthcare) at room temperature (25 °C). Equal concentrations of gp59, gp60, gp61 and gp68 were immobilized on channels of the carboxymethyldextran-modified (CM5) sensor chip to about 280 response units (RU). To collect data for single-cycle kinetic analysis, a single cycle of consecutive injections of diverse signaling molecules at increasing concentrations (6.25 µM, 12.5 µM, 25 µM, 50 µM and 100 µM) was performed. The analytes were diluted in binding buffer (20 mM HEPES pH 7.5, 200 mM NaCl, and 0.05% (v/v) Tween-20) and injected over the chip at a flow rate of 30 µl min⁻¹. Each concentration was injected for 60 s without intermediate regeneration, followed by a final dissociation phase of 600 s. Data was fit with a model describing a bivalent analyte. Kinetic rate constants were extracted from this curve fit using Biacore evaluation software (GE Healthcare).

### HPLC analysis

For analysis of ligand sequestering, a reaction mixture containing 50 μM 2′,3′-cAA and 100 μM gp59, was incubated in 10 mM Tris-Hcl pH 8.0 and 500 mM NaCl at 16°C for 30 min. The samples were filtered through a 3 kDa cutoff ultrafiltration tube, and the flow-through fraction was analyzed using an InfinityLab Poroshell 120 AQ-C18 column (4.6 × 150 mm, 2.7 μm; Agilent). Chromatographic separation was performed at a flow rate of 0.4 ml/min under isocratic elution with 97% 20 mM potassium phosphate buffer (pH 6.0) and 3% acetonitrile. Detection was carried out at 254 nm. To examine the release of 2′,3′-cAA, gp59 pre-bound with 2′,3′-cAA was treated with SDS to a final concentration of 1% to inactivate the protein. The mixture was then filtered through a 0.22 µm filter and subsequently subjected to HPLC analysis to detect the released 2′,3′-cAA.

For analysis of ligand sequestering, reaction mixtures containing 100 μM 1′′,3′-gcADPR and 200 μM gp68 (or its truncated variant) were incubated in 10 mM Tris-HCl pH 8.0 and 200 mM NaCl at 16°C for 60 min. The samples were filtered through a 3 kDa cutoff ultrafiltration tube, and the flow-through fractions were analyzed using an InfinityLab Poroshell 120 AQ-C18 column (4.6 × 150 mm, 2.7 μm; Agilent). Chromatographic separation was performed at a flow rate of 0.4 ml/min under isocratic elution with 97% 20 mM potassium phosphate buffer (pH 6.0) and 3% acetonitrile. Detection was carried out at 254 nm. To examine the release of 1′′,3′-gcADPR, gp68 (or its truncated variants) pre-bound with the respective ligand was treated with SDS to a final concentration of 1% to inactivate the protein. The mixture was then filtered through a 0.22 µm filter and subsequently subjected to HPLC analysis to detect the released 1′′,3′-gcADPR.

### SEC-MALS (Size Exclusion Chromatography-Multi-Angle Light Scattering)

SEC-MALS measurements were performed using a protein sample at a concentration of 6 mg/mL with a total volume of 100 μL. Prior to analysis, both the protein sample and the corresponding buffer were filtered through a 0.22 μm membrane filter to remove dust particles and aggregates. The instrument and temperature-control system were equilibrated, and the laser source was preheated for at least 30 min before data acquisition. Filtered buffer was first loaded into a clean cuvette to record the solvent background signal. Subsequently, the protein sample was added to the cuvette and allowed to equilibrate to the measurement temperature before scattering intensity data were collected at the designated detection angle.

## References

1. Cohen, D. et al. Cyclic GMP-AMP signalling protects bacteria against viral infection. Nature 574, 691–695 (2019).

2. Ofir, G. et al. Antiviral activity of bacterial TIR domains via immune signalling molecules. Nature 600, 116–120 (2021).

3. Ceyssens, P.-J. et al. Molecular and physiological analysis of three Pseudomonas aeruginosa phages belonging to the ‘N4-like viruses’. Virology 405, 26–30 (2010).

4. Georjon, H. & Bernheim, A. The highly diverse antiphage defence systems of bacteria. Nat. Rev. Microbiol. 21, 686–700 (2023).

5. Hochhauser, D. & Sorek, R. Manipulation of the nucleotide pool in human, bacterial and plant immunity. Nat. Rev. Immunol. 26, 7–22 (2026).

6. Millman, A., Melamed, S., Amitai, G. & Sorek, R. Diversity and classification of cyclic-oligonucleotide-based anti-phage signalling systems. Nat Microbiol 5, 1608–1615 (2020).

7. Athukoralage, J. S. & White, M. F. Cyclic nucleotide signaling in phage defense and counter-defense. Annu. Rev. Virol. 9, 451–468 (2022).

8. Duncan-Lowey, B., McNamara-Bordewick, N. K., Tal, N., Sorek, R. & Kranzusch, P. J. Effector-mediated membrane disruption controls cell death in CBASS antiphage defense. Mol. Cell 81, 5039–5051.e5 (2021).

9. Lau, R. K. et al. Structure and Mechanism of a Cyclic Trinucleotide-Activated Bacterial Endonuclease Mediating Bacteriophage Immunity. Mol. Cell 77, 723–733.e6 (2020).

10. Huiting, E., et al. CBASS limits bacteriophage production while maintaining cell viability in *Pseudomonas aeruginosa*. *bioRxiv* (2026) doi:10.64898/2026.02.24.707611.

11. Whiteley, A. T. et al. Bacterial cGAS-like enzymes synthesize diverse nucleotide signals. Nature 567, 194–199 (2019).

12. Sabonis, D. et al. TIR domains produce histidine-ADPR as an immune signal in bacteria. Nature (2025) doi:10.1038/s41586-025-08930-2.

13. Leavitt, A. et al. Viruses inhibit TIR gcADPR signalling to overcome bacterial defence. Nature 611, 326–331 (2022).

14. Shi, Y. et al. Structural characterization of macro domain-containing Thoeris antiphage defense systems. Sci. Adv. 10, eadn3310 (2024).

15. Rousset, F. et al. TIR signaling activates caspase-like immunity in bacteria. Science 387, 510–516 (2025).

16. Roberts, C. G. et al. Bacterial TIR-based immune systems sense phage capsids to initiate defense. Nat. Microbiol. 10, 2892–2902 (2025).

17. Wang, L., Zheng, R. & Zhang, L. Sequestering survival: sponge-like proteins in phage evasion of bacterial immune defenses. Front. Immunol. 16, 1545308 (2025).

18. Huiting, E. et al. Bacteriophages inhibit and evade cGAS-like immune function in bacteria. Cell 186, 864–876.e21 (2023).

19. Jenson, J. M., Li, T., Du, F., Ea, C.-K. & Chen, Z. J. Ubiquitin-like conjugation by bacterial cGAS enhances anti-phage defence. Nature (2023) doi:10.1038/s41586-023-05862-7.

20. Tal, N. et al. Structural modeling reveals phage proteins that manipulate bacterial immune signaling. Science 391, eaea1761 (2026).

21. Li, D. et al. Single phage proteins sequester signals from TIR and cGAS-like enzymes. Nature 635, 719–727 (2024).

22. Cao, X. et al. Phage anti-CBASS protein simultaneously sequesters cyclic trinucleotides and dinucleotides. Mol. Cell 84, 375–385.e7 (2024).

23. Hadary, R., et al. Functional diversity of phage sponge proteins that sequester host immune signals. *bioRxivorg* (2025) doi:10.1101/2025.08.24.671296.

24. Sullivan, A. E. et al. The Panoptes system uses decoy cyclic nucleotides to defend against phage. Nature 647, 988–996 (2025).

25. Doherty, E. E. et al. A miniature CRISPR-Cas10 enzyme confers immunity by inhibitory signalling. Nature 647, 997–1004 (2025).

26. Wittmann, J. et al. From orphan phage to a proposed new family-the diversity of N4-like viruses. Antibiotics (Basel) 9, E663 (2020).

27. Ferry, T. et al. Personalized bacteriophage therapy to treat pandrug-resistant spinal Pseudomonas aeruginosa infection. Nat. Commun. 13, 4239 (2022).

28. Zang, Z. et al. Chemical inhibition of a bacterial immune system. Cell Host Microbe 34, 263–277.e11 (2026).

29. Lebreton, F., et al. A panel of diverse Pseudomonas aeruginosa clinical isolates for research and development. JAC Antimicrob. Resist. 3, dlab179 (2021).

30. Zang, Z., et al. Chemical inhibition of a bacterial immune system 1. *bioRxivorg* (2025) doi:10.1101/2025.02.20.638879.

31. Jurczak-Kurek, A. et al. Biodiversity of bacteriophages: morphological and biological properties of a large group of phages isolated from urban sewage. Sci. Rep. 6, 34338 (2016).

32. Gilchrist, C. L. M. & Chooi, Y.-H. Clinker & clustermap.Js: Automatic generation of gene cluster comparison figures. Bioinformatics 37, 2473–2475 (2021).

33. Csörgő, B. et al. A compact Cascade-Cas3 system for targeted genome engineering. Nat. Methods 17, 1183–1190 (2020).

34. Bartual, S. G., Garcia-Doval, C., Alonso, J., Schoehn, G. & van Raaij, M. J. Two-chaperone assisted soluble expression and purification of the bacteriophage T4 long tail fibre protein gp37. Protein Expr. Purif. 70, 116–121 (2010).

35. Marti, R. et al. Long tail fibres of the novel broad-host-range T-even bacteriophage S16 specifically recognize Salmonella OmpC. Mol. Microbiol. 87, 818–834 (2013).

36. Schulz, E. C. et al. Crystal structure of an intramolecular chaperone mediating triple-beta-helix folding. Nat. Struct. Mol. Biol. 17, 210–215 (2010).

37. North, O. I. & Davidson, A. R. Phage proteins required for tail fiber assembly also bind specifically to the surface of host bacterial strains. J. Bacteriol. 203, e00406–20 (2021).

38. Saha, S. et al. F-type pyocins are diverse noncontractile phage tail-like weapons for killing Pseudomonas aeruginosa. J. Bacteriol. 205, e0002923 (2023).

39. Chang, R. B. et al. A widespread family of viral sponge proteins reveals specific inhibition of nucleotide signals in anti-phage defense. Mol. Cell 85, 3151–3165.e6 (2025).

40. Tesson, F. et al. Exploring the diversity of anti-defense systems across prokaryotes, phages and mobile genetic elements. Nucleic Acids Res. 53, gkae1171 (2025).

41. Pinilla-Redondo, R. et al. Discovery of multiple anti-CRISPRs highlights anti-defense gene clustering in mobile genetic elements. Nat. Commun. 11, 5652 (2020).

42. Samuel, B., Mittelman, K., Croitoru, S. Y., Ben Haim, M. & Burstein, D. Diverse anti-defence systems are encoded in the leading region of plasmids. Nature 635, 186–192 (2024).

43. Jha, N. et al. Gaia: An AI-enabled genomic context-aware platform for protein sequence annotation. Sci. Adv. 11, eadv5109 (2025).

44. Yuchi, Z. et al. Crystal structures of ryanodine receptor SPRY1 and tandem-repeat domains reveal a critical FKBP12 binding determinant. Nat. Commun. 6, 7947 (2015).

45. Zalk, R. et al. Structure of a mammalian ryanodine receptor. Nature 517, 44–49 (2015).

46. Melville, Z. et al. A drug and ATP binding site in type 1 ryanodine receptor. Structure 30, 1025–1034.e4 (2022).

47. Holm, L., Laiho, A., Törönen, P. & Salgado, M. DALI shines a light on remote homologs: One hundred discoveries. Protein Sci. 32, e4519 (2023).

48. Gao, L. A. et al. Prokaryotic innate immunity through pattern recognition of conserved viral proteins. Science 377, eabm4096 (2022).

49. Lee, H., et al. Diverse bacterial pattern recognition receptors sense the conserved phage proteome. *bioRxivorg* (2026) doi:10.64898/2026.01.04.697583.

50. Pawluk, A., Bondy-Denomy, J., Cheung, V. H. W., Maxwell, K. L. & Davidson, A. R. A new group of phage anti-CRISPR genes inhibits the type I-E CRISPR-Cas system of Pseudomonas aeruginosa. MBio 5, e00896 (2014).

51. Rauch, B. J. et al. Inhibition of CRISPR-Cas9 with Bacteriophage Proteins. Cell 168, 150–158.e10 (2017).

52. Marino, N. D. et al. Discovery of widespread type I and type V CRISPR-Cas inhibitors. Science 362, 240–242 (2018).

53. Choi, K.-H. & Schweizer, H. P. mini-Tn7 insertion in bacteria with single attTn7 sites: example Pseudomonas aeruginosa. Nat. Protoc. 1, 153–161 (2006).

54. Li, H. & Durbin, R. Fast and accurate short read alignment with Burrows-Wheeler transform. Bioinformatics 25, 1754–1760 (2009).

55. Robinson, J. T. et al. Integrative genomics viewer. Nat. Biotechnol. 29, 24–26 (2011).

56. Abramson, J. et al. Accurate structure prediction of biomolecular interactions with AlphaFold 3. Nature 630, 493–500 (2024).

57. Emsley, P., Lohkamp, B., Scott, W. G. & Cowtan, K. Features and development of coot. Acta Crystallogr. D Biol. Crystallogr. 66, 486–501 (2010).

58. Adams, P. D. et al. PHENIX: building new software for automated crystallographic structure determination. Acta Crystallogr. D Biol. Crystallogr. 58, 1948–1954 (2002).

